# Intrinsic polarization cues interfere with pheromone gradient sensing in *S. cerevisiae*

**DOI:** 10.1101/502856

**Authors:** Gustavo Vasen, Alejandro Colman-Lerner

## Abstract

Polarity decisions are central to many processes, including mitosis and chemotropism. In *S. cerevisiae*, budding and mating projection (MP) formation use an overlapping system of cortical landmarks that converge on the small G-protein Cdc42. However, pheromone gradient sensing must override the Rsr1-dependent internal polarity cues used for budding. Using this model system, we asked what happens when intrinsic and extrinsic spatial cues are misaligned. Is there competition, or collaboration? By live cell microscopy and microfluidics technics we uncovered three previously overlooked features of this signaling system. First, the cytokinesis-associated polarization patch serves as a polarity landmark independently of all known cues. Second, the Rax1-Rax2 complex functions as novel pheromone promoted polarity cue in the distal pole of the cells. Finally, we showed that internal cues remain active during pheromone gradient tracking and that they interfere with this process biasing the location of MPs, since yeast defective in internal cue utilization align significantly better than wild-type with artificially generated pheromone gradients.

## Introduction

Cell polarity is central to all living organisms for proliferation, differentiation and morphogenesis. Examples of polarity associated processes are widely distributed during embryo development, chemotaxis of bacteria or immune cells, chemotropism observed during axon guidance, cell migration, epithelial polarity and asymmetric cell division. Similarly, malfunction of polarity mechanisms is associated with diverse pathological conditions such as neurodegenerative diseases, genetic disorders and cancer (Aguilar et al., 2017; Egorov and Polishchuk, 2017; Haga and Ridley, 2016).

The small G protein Cdc42, discovered in yeast, is the master regulator of cell polarity throughout eukaryotic cells (Adams et al., 1990; Etienne-Manneville, 2004). During polarity establishment, cells are able to interpret both extracellular cues and intrinsic cell landmarks to generate intracellular asymmetry (Yogev and Shen, 2017). However, a question that still remains unanswered is how the many signaling pathways that converge on Cdc42 are interpreted, processed and integrated so that the cell makes an appropriate response. This is especially interesting in the cases where input cues are conflicting.

To address this issue, we focused on the Cdc42 polarization system in *S. cerevisiae*, which is arguably the best studied example of eukaryotic cell polarity (Goryachev and Leda, 2017; Park and Bi, 2007). Yeast polarize their growth during cell division to form a bud and in response to sexual pheromone gradients to find a mate. In both processes, a core set of proteins concentrate to form a polarity patch. Cdc42 is locally activated (exchanging GDP for GTP) by its GEF Cdc24 and this interaction is stabilized by the scaffold protein Bem1 (Butty et al., 2002; Zheng et al., 1994). In a positive feedback loop, Cdc42-GTP recruits more of the Bem1-Cdc24 complex, helping to activate more Cdc42 (Kozubowski et al., 2008). Therefore, polarization in this system is a highly self-reinforcing process. Cdc42-GTP also recruits the formins Bni1 and Bnr1, which nucleate linear actin filaments that direct vesicle delivery into the polarity patch. Bni1 is part of the polarisome, a complex organized by the scaffold protein Spa2, that acts as the focal point for polymerization of actin monomers into actin cables (Sheu et al., 1998). The concerted action of these proteins allows polarized cell growth.

The polarization required for bud formation is directed to specific sites of the cell surface through landmark proteins, known as budding cues. During mating, haploid cells of the mating types MATa and MATα secrete a and α-factor pheromones, respectively, forming gradients that the cell of the opposite mating type detects and tracks (Jackson and Hartwell, 1990). In doing so, they form mating projections (MP) that polarize cell growth in the direction of the partner. Budding and MP formation share the core polarity machinery based on Cdc42. Moreover, it has been shown that in the absence of pheromone gradients, i.e. isotropic pheromone stimulation, budding cues remain active and can direct the position of MPs (Madden and Snyder, 1992). This posits a conflicting scenario where intrinsic cues can compete with external polarity signals and it provides a good platform to study the integration of spatial information by the Cdc42 module in eukaryotes.

Budding cues are localized at either pole of the cell (Figure 1). The proximal pole is defined by the place where cytokinesis occurred and coincides with the location of the birth scar, a characteristic thinning in the cell wall caused by the action of daughter-specific chitinases (Colman-Lerner et al., 2001). On the mother side, the separation of each daughter cell leaves a chitin ring structure known as bud scar. The distal pole is located opposite to the birth scar along the major axis of the cell. Wild-type (WT) haploid cells follow the *axial* budding pattern where buds are formed exclusively at the proximal pole. In contrast, diploid cells use a *bipolar* pattern placing the first bud predominantly in the distal pole and, in subsequent divisions, at either pole (Chant and Pringle, 1995; Freifelder, 1960). At the molecular level, Axl2/Bud10 and Bud9 are the proximal cues used by haploid and diploid cells, respectively, while Bud8 is the distal landmark that guides budding exclusively in diploid cells (Bi and Park, 2012). In late G1, these protein landmarks locally activate the small G-protein Rsr1/Bud1 through the recruitment of its GEF Bud5 (Kang et al., 2001; Krappmann et al., 2007). Active Rsr1-GTP brings Cdc24 to the membrane which actives Cdc42-GTP to initiate budding (Zheng et al., 1995). In the absence of Rsr1, diploid and haploid cells bud randomly (Bender and Pringle, 1989; Chant and Herskowitz, 1991). Interestingly, in haploid cells both axial and bipolar landmark proteins are expressed but if Axl2 is functional, it dominates over Bud8 or Bud9, presumably because of a higher ability to recruit Bud5.

**Figure 1.**
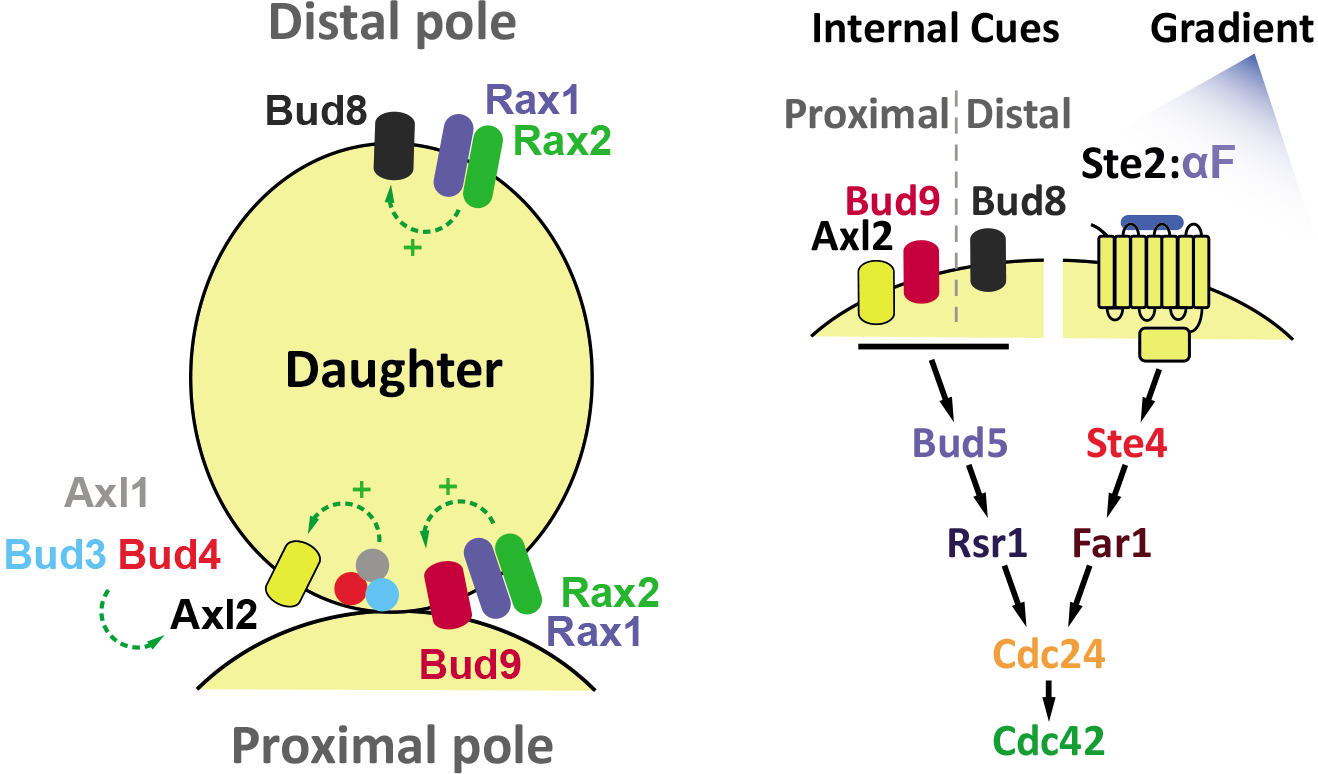
Establishment of bud site selection program and cell polarity. **Left.** The proximal and distal pole of a daughter cell is specified relative to the position of the mother cell. For budding haploid cells utilize Axl2, the axial landmark, which is positioned in every cell cycle at the proximal pole by the action of Bud3, Bud4 and the haploid-specific protein Axl1. In the absence of Axl2, diploid cells use bipolar landmarks, distal Bud8 and proximal Bud9, to choose the budding site. The Rax1-Rax2 complex participates in the localization of both Bud8 and Bud9. **Right.** In late G1, Axl2, Bud8 or Bud9, depending on cell type and condition, recruit Bud5 where it activates Rsr1 which, in turn, brings Cdc24, the Cdc42’s GEF, to the membrane to generate a concentrated pool of active Cdc42-GTP. During mating, in MATa cells, α-factor binds to its receptor Ste2 and the activated receptor dissociates the G-protein trimer into G and Gβ/Ste4. Ste4 recruits the adaptor protein Far1 in complex with Cdc24 to the side of the membrane exposed to the higher concentration of α-factor. As in the case of budding, Cdc24 locally activates Cdc42.

It has been suggested that under isotropic pheromone stimulation haploid cells form MPs at the proximal pole, guided by the budding cues (Madden and Snyder, 1992). The original (and nearly exclusive) evidence supporting this notion comes from the observation that in strains with random budding patterns, such as Δ*rsr1*, MPs are also randomly placed. However, the positions of MPs were measured relative to a bud scar that was randomly placed. Then, no definitive positional information about MPs can be extracted from data of this mutant. Thus, the question remains as to whether MPs follow budding cues.

Binding of pheromone to a G-protein Coupled Receptor (Ste2 in MATa and Ste3 in MATα) yeast activates the pheromone response system, which is nearly identical in both sexes (Figure 1). Upon α-factor binding, active Ste2 causes the dissociation of the Ste4/Gβ-Ste18/Gγ dimer from Gpa1/Gα (Bardwell, 2004). Free Gβγ recruits the scaffold protein Ste5 and the adaptor protein Far1 to the plasma membrane. In turn, Ste5 activates a MAPK-dependent signaling cascade that induces mating related genes, stimulates polarization and arrests the cell cycle in G1 phase (Dohlman and Thorner, 2001; Pryciak and Huntress, 1998). Far1 associates with Cdc24 and functions as the link between ligand-bound active receptors and the Cdc42 polarization module (Butty et al., 1998; Nern and Arkowitz, 1999). In this way the pheromone gradient biases the cytoskeleton and cell growth in the direction of the mating partner. If Ste4 and the polarity machinery are decoupled (like in strains with *FAR1* alleles that do not bind Gβ or the *cdc24-m1* allele that does not associate with Far1) MPs cannot track gradients and form next to the last bud scar (Nern and Arkowitz, 1998; Valtz et al., 1995).

The interaction between budding cues and pheromone gradient tracking is still controversial. Two main ideas are thought to govern gradient detection in *S. cerevisiae*: global and local sensing (Hegemann and Peter, 2017). In the first case, polarization would form directly towards the pheromone source as a global reaction of the system. For that to take place, it was thought that proper detection of pheromone gradients involved inactivation of (Roemer et al., 1996) or interference with, internal budding cues. The latter may be accomplished, for example, by biasing Cdc24 association towards Far1 instead of Rsr1 (Nern and Arkowitz, 1999). In the local sensing model, an “exploratory” polarity patch, established independently of gradient direction, would correct its position through local sampling. This is possible because the polarity patch is highly dynamic and mobile, especially at low α-factor concentrations (Dyer et al., 2013; Nern and Arkowitz, 2000). In this scenario, inactivation of budding cues may not be necessary. In fact, it was recently suggested that during gradient sensing, the site of polarity establishment starts at budding cues independently of gradient direction. Later on, the polar cap location is corrected towards the pheromone source (Hegemann et al., 2015). To account for this correction process many mechanisms where proposed: an actin-driven perturbation caused by vesicle fusion (Dyer et al., 2013), a differential internalization of the pheromone receptor as a result of its asymmetric phosphorylation (Ismael et al., 2016), and a pheromone dose-dependent local regulation of Cdc42 activity through the nuclear sequestration of Cdc24 caused by the MAPK Fus3 (Hegemann et al., 2015). Each of these three proposed mechanisms may account for a greater polar cap mobility at low pheromone concentrations (around the K_D_ of the α-factor-Ste2 interaction), which might explain why gradient detection in this region is highest. In the actin-dependent mechanism, vesicles delivering unoccupied receptors would first dilute polarity factors in the patch at low doses, since binding of α-factor is slow. In the Fus3-dependent case, Fus3 activity is proposed to confer stability to the polarity patch, hence restricting its mobility as a function of the pheromone dose.

Considering that budding cues remain active during pheromone stimulation, we decided to revisit this simple, yet unanswered question: what happens when intrinsic and extrinsic spatial cues are misaligned? Is there competition, so that internal cues interfere with efficient gradient sensing, or collaboration? While in the process of answering this question, we discovered that the default polarization site in response to pheromone is the result of a much more complex system of landmarks, which has remained, surprisingly, unnoticed. Once uncovered, we found that, indeed, intrinsic polarity sites can interfere with pheromone gradient sensing.

## Results

### Default polarization sites in response to α-factor

To answer the question of how internal and external cues interact during pheromone gradient sensing, we first revisited the role of internal cues during isotropic stimulation. Classic work showed that in this situation yeast use a default mechanism, positioning their MPs not randomly but employing the same internal polarization cues used during budding (Madden and Snyder, 1992). These landmarks are located in both poles of the cell: the proximal and the distal pole, relative to the location of the mother. To confirm these observations, we studied the default position of MPs in our wild-type strain derived from the W303 lineage. We focused on daughter cells because their poles are easily recognized by live cell imaging. First, we defined the default budding site by measuring the angle between the first bud of a daughter cell and the proximal pole (i.e. the position of its mother) (Figure 2A). Around 30% of the population budded in the proximal pole with angles smaller than 60° and almost 60% of the cells used the distal pole (angles>120°). This corresponds to a mixture of axial and bipolar budding patterns instead of the purely proximal budding expected for haploid cells. The abnormal bud site selection of W303 is known and it is thought to be due to a defective positioning of the axial landmark Axl2 (see below). When exposed to uniform saturating concentrations of α-factor, daughter cells also formed MPs at both poles. However, the fraction of cells that chose the proximal and the distal poles was different than during budding (Figure 2A), suggesting the existence of extra regulation for the selection of MPs. By analyzing the behavior of different cells, it became evident that daughter cells that were buds at the time of pheromone stimulation, and which therefore make an MP only after finishing the cytokinesis, had a strong tendency to form MPs at the proximal pole. Thus, we decided to classify daughter cells in 2 categories (Figure 2B): cells born before (BB) α-factor exposure, which were already in G1 and could execute a chemotropic response readily and cells born after (BA) pheromone stimulation, which would form MPs only after entering the new G1 phase. This sorting into BA and BB cells turned out to be fundamental for the understanding of the relationship between budding cues and MP default site selection. By analyzing the response to different α-factor concentrations (Figure 2C), we observed clear differences in the behavior of BA and BB cells. Cells already in G1 (BB), despite budding at either pole of the cell (Figure 2A), formed MPs almost exclusively in the distal pole, at all pheromone concentrations tested. On the other hand, BA cells showed a dose-dependent site selection: at low α-factor concentrations, cells polarized in the distal pole, as BB cells, but at higher pheromone doses BA cells formed MPs in the proximal pole. Hence, the behavior shown in Figure 2A, obtained at a saturating pheromone concentration, was the result of summing proximal pole-choosing BA and distal pole-choosing BB daughters. Given this scenario, in what follows we searched for the molecular landmarks that define the position of MPs in the proximal and the distal poles and for the mechanism responsible for the dose-dependent switch in the choice of location of MPs in BA cells.

**Figure 2.**
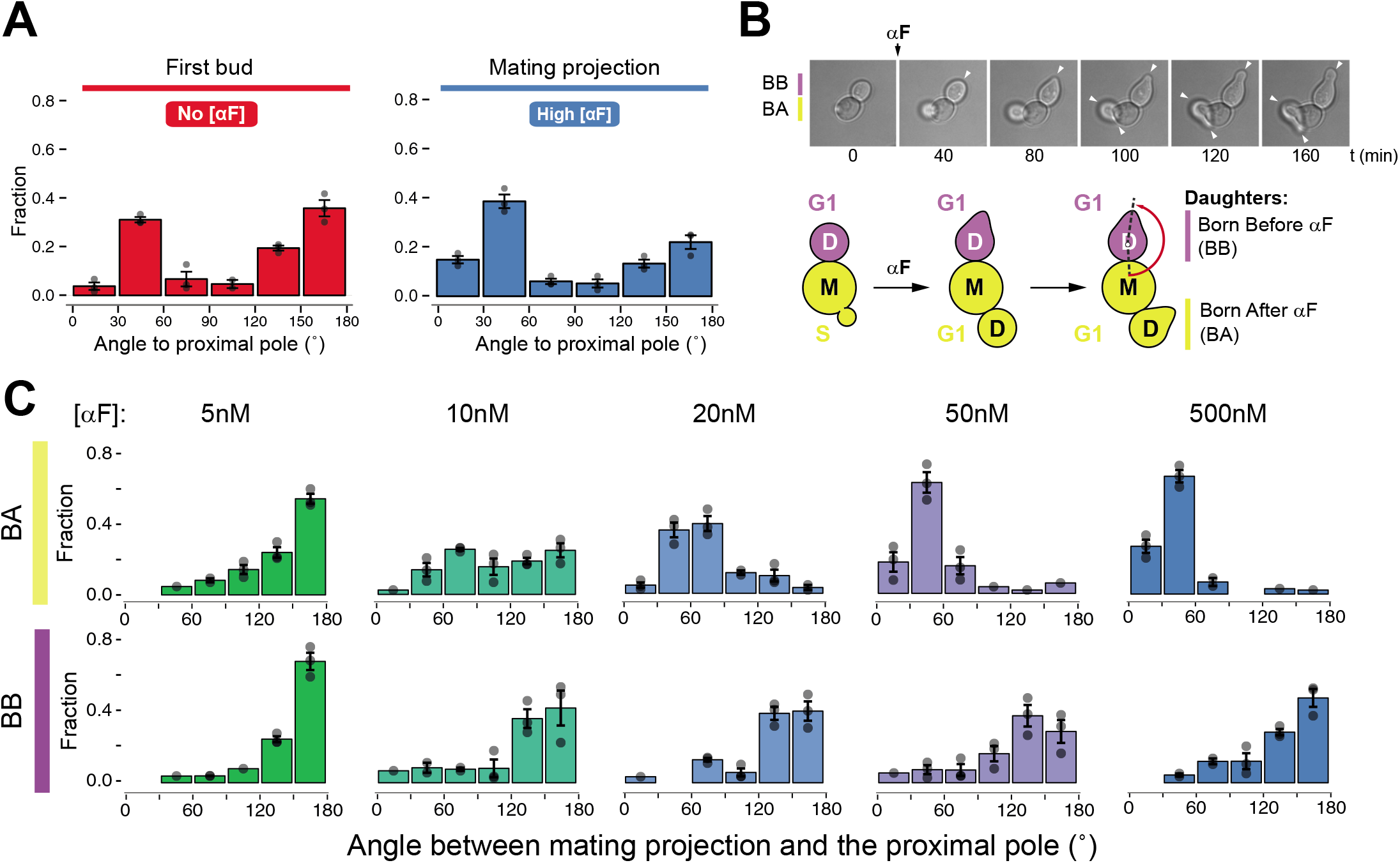
Mating projection default position in W303. **A.** Exponential cultures of W303 Δ*bar1* strain (ACL379) were subjected to live cell imaging in the absence (No [αF]) or presence of 1μM α-factor (High [αF]). The angle of the first bud or the mating projection (MP) was measured in daughter cells relative to the proximal pole (i.e. where the mother cell is localized). In this and subsequent panels, the fraction of cells in each 30°-bin was plotted for at least 3 independent experiments. The bar and error bars represent the mean ± SEM and the points reflect the data from each assay. **B.** Top. Time-lapse images of cells stimulated with 1μM αF. Arrowheads indicate the position of the mating projection. Bottom. Classification into 2 categories of daughter cells used throughout the paper: born before stimulation (BB) or born after stimulation (BA). **C.** MP angle distribution relative to the proximal pole of BA and BB cells in W303 Δ*bar1* yeast (ACL379) exposed to different pheromone doses.

Since W303 has an abnormal bud site selection, first we tested whether the behaviors outlined above were a consequence of W303 having a weak Axl2 axial landmark. This is due to the inactivating mutation in *BUD4* that is present in this strain background (Ralser et al., 2012; Voth et al., 2005). Bud4 links Axl2 to the septin ring at the cytokinesis site and helps, in complex with Bud3 and Axl1, setting the proximal cue (Figure 1). Indeed, S288C, which bears a wild-type *BUD4* allele, budded exclusively at the proximal pole and formed MPs preferentially in the same site. However, at low pheromone concentrations a significant fraction of S288C cells (20-30%) selected the distal pole for MPs (Figure 3B). We then constructed two strains in which we swapped *BUD4* alleles between backgrounds resulting in a W303 *BUD4*^*S288C*^ and an S228C *bud4*^*W303*^ (Figure 3). The S228C *bud4*^*W303*^ exhibited the exact bud and MPs site selection we observed in W303 (Figure 3C). Conversely, in W303, the expression of *BUD4* reestablished the axial budding but, notably, only partially changed the default site for MPs (Figure 3D). At saturating α-factor concentrations, BB cells formed projections mostly in the proximal pole but at low pheromone doses in BA cells there was a clear bias towards making distal MPs, suggesting that W303 has stronger mechanisms than S288C to position polarizations in the distal pole besides the Bud4-Axl2 axis.

**Figure 3.**
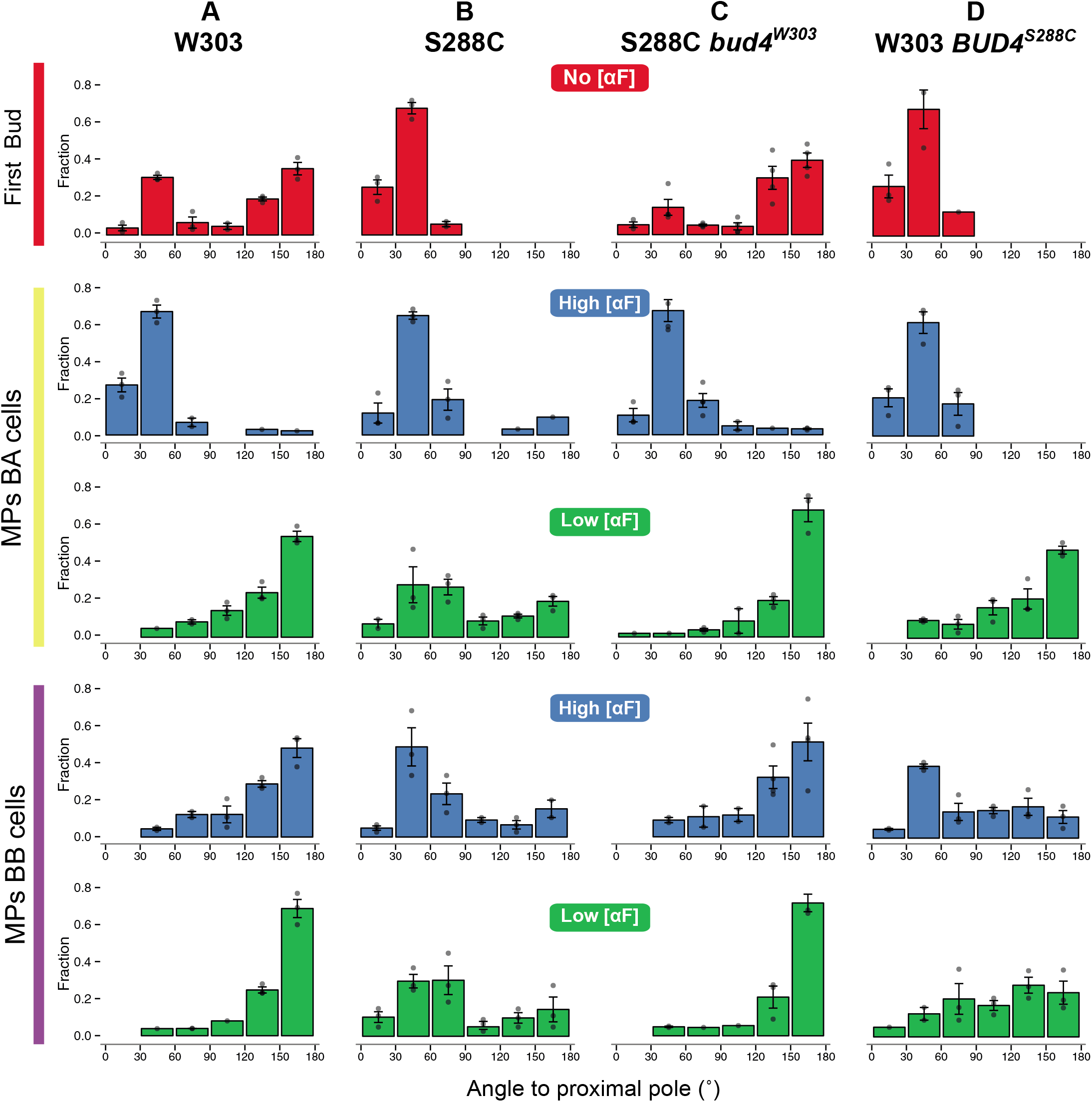
Influence of BUD4 on mating projection positioning. Distribution of angles of first bud or mating projections (MPs) relative to the proximal pole in *bud4* or *BUD4* strains. Different panels correspond to angles of first buds (No [αF]), 1μM (High [αF]) or 5nM (Low [αF]) pheromone concentrations in both BA and BB cells. **A**. W303 Δ*bar1* strain (ACL379). **B**. S288C BY4741 Δ*bar1* (YDV5654). **C**. S288C BY4741 Δ*bar1 bud4*^*W303*^ (YGV5831). **D.** W303 Δbar1 *BUD4*^*S288C*^ cells (YGV5765).

The differences between BA and BB daughters were unexpected. Thus, we studied the dynamics of the polarity patch in these two classes of daughters using Bem1 fused with 3 tandem copies of mNeonGreen (Bem1-3xmNG) in W303 (Ventura et al., 2014). In cycling cells, Bem1-3xmNG was in the cytoplasm in G1 cells and translocated to the presumptive bud location after Start, as previously published (Gulli et al., 2000). In G2/M, Bem1-3xmNG was restricted to the periphery of the bud and finally it relocated to mother-daughter neck in late anaphase, where it stayed during cytokinesis (Figure 4A). Based on the localization of Bem1-3xmNG at the time of α-factor exposure, we measured the angle of MPs in 4 different groups: *incipient bud*, *bud periphery*, *bud neck* and *cytoplasmic*. BB cells correspond to daughter cells from the last group, in which MPs formed by *de novo* polarization in the distal pole (Figure 4B and 3C). The remaining groups comprised BA cells (Figure 4B). We observed that the location of MPs in BA cells was not homogeneous but it depended on how far along in the cell cycle each cell was at the time of stimulation with α-factor. In the *incipient bud* group, MPs were overwhelmingly proximal. In the *bud periphery* group, they were less so, with some cells choosing larger angles. Finally, in the last group, in which Bem1 was already at the bud neck, the MP distribution centered between 60° and 90°. Detailed observation of Bem1-3xmNG in BA cells showed that stimulation with pheromone prevented the dispersal of the polarity patch that normally occurs when cells enter G1 (Figure 4C). Instead, the polarity patch involved in cytokinesis persists, and it is used directly for MP polarization. This is possible since we detected signaling proteins from the pheromone pathway like Ste4/Gβ localized in the neck during cytokinesis even in the absence of α-factor (Figure 4D). When pheromone was added to these cells, the Ste4 patch remained in the membrane, slightly moving to the side of the cytokinesis site (Figure 4D, bottom).

**Figure 4.**
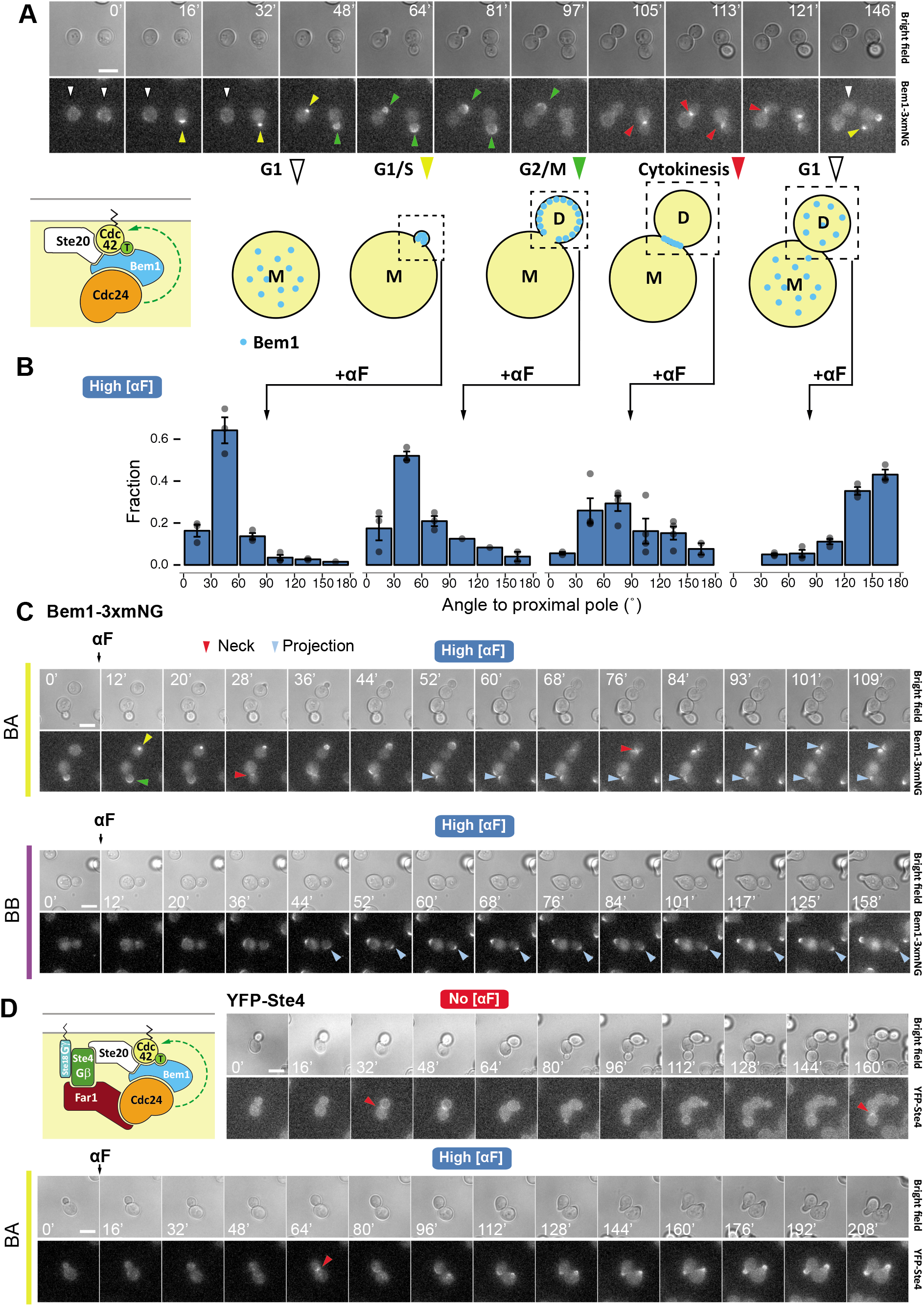
Polarity patch dynamics in default mating projection site selection. **A.** Fluorescence microscopy time-lapse of yeast expressing Bem1 fused to three mNeonGreen fluorescent proteins (Bem1-3xmNG, YGV5097) through the cell cycle. The diagram below summarizes the localization of Bem1. **B.** Daughter cells were divided in 4 groups based on the localization of Bem1-3xmNG at the time of pheromone stimulation (1 μM). The angle between mating projection and the proximal pole was measured. The fraction of cells in each 30°-bin was plotted for at least 3 independent experiments. The bar and error bars represent the mean ± SEM and the points reflect the data from each assay. **C.** Bem1-3xmNG dynamics in BA and BB cells exposed to 1μM α-factor. Arrowheads denote the different localizations of Bem1-3xmNG as represented in the diagram above. **D.** Time lapse of N-terminally YFP tagged Ste4 (YFPSte4, TCY3064) in cycling cells (No [αF]) or cells exposed to 1 μM (High [αF]). For all images, times of acquisition is indicated in minutes and scale bar represents 5μm.

We interpret the differences between the three groups of BA cells assuming the existence of a mechanism activated around late anaphase that later prevents MP formation at the site of cytokinesis. When α-factor is added before this inhibitory mechanism is in place, the pheromone signaling blocks it, and the MP forms proximal. Otherwise, the polar cap moves away from the neck before it forms the MP. In what follows, unless specifically stated, we excluded the bud neck BA group from the analysis for technical reasons (see Methods).

### BA cells use the cytokinesis patch to initiate an MP, independently of budding cues

The above observations led us to hypothesize that the proximal pole was chosen at saturating pheromone in BA cells (*incipient bud* and *bud periphery* groups) not because of the action of axial budding cues but due to the presence of a pre-polarized structure at the neck. At lower α-factor doses, as the polarity patch mobility increases (Dyer et al., 2013), the polar cap may “wander” off from the neck towards the distal pole (Figure 2C and Figure 5A). Indeed, the polarity patch of Bem1-3xmNG in BA cells exposed to 5nM α-factor migrates away from the neck (Figure 5B). To test the idea that patch wandering plays a role in the switch from proximal (at high αF) to distal (at low αF) MP localization, we artificially increased the polarity patch mobility by two different genetic strategies. First, we destabilized the polarisome, the molecular structure that maintains vesicle delivery focused at the polarity site, by deleting *SPA2*. This mutation causes a broadening of the MPs (peanut phenotype (Valtz and Herskowitz, 1996) at high α-factor concentrations) and, similar to the effect of lowering the dose of pheromone, it resulted in reduced MPs at the proximal pole (Figure 5A). In agreement with this, the Bem1-3xmNG cap in *Δspa2* cells had elevated mobility even at a high α-factor concentration (Figure 5B). This supported the notion that movement of the polarity patch allows it to escape the neck prepolarization. At saturating α-factor concentrations in Δ*spa2*, MPs are formed away from the proximal pole at various angles suggesting that the polarity patch still has restricted mobility. Further increasing patch mobility in Δ*spa2* by lowering the concentrations of pheromone resulted in MPs forming at the distal pole (Figure 5A). The second approach consisted in increasing the polarity patch wandering by expressing a mutant receptor allele (*STE2-F204S)* that does not bind ligand (Dosil et al., 1998) in cells that also express (WT) Ste2. As we have previously showed, this mutant receptor actively inhibits pheromone signaling by its association with Sst2, the Gα/Gpa1 GAP (Bush et al., 2016). Cells that co-express wild-type and mutant *STE2* form twisted MPs likely due to the higher mobility of the polarity patch (Figure 5A inset). In this case, Bem1-3xmNG patch moved out from the neck and MPs were formed elsewhere (Figure 5A and B). These results support the idea that proximal MPs observed in BA cells at high pheromone concentrations are the combined effect of low polarity patch mobility and a pre-polarized starting state.

**Figure 5.**
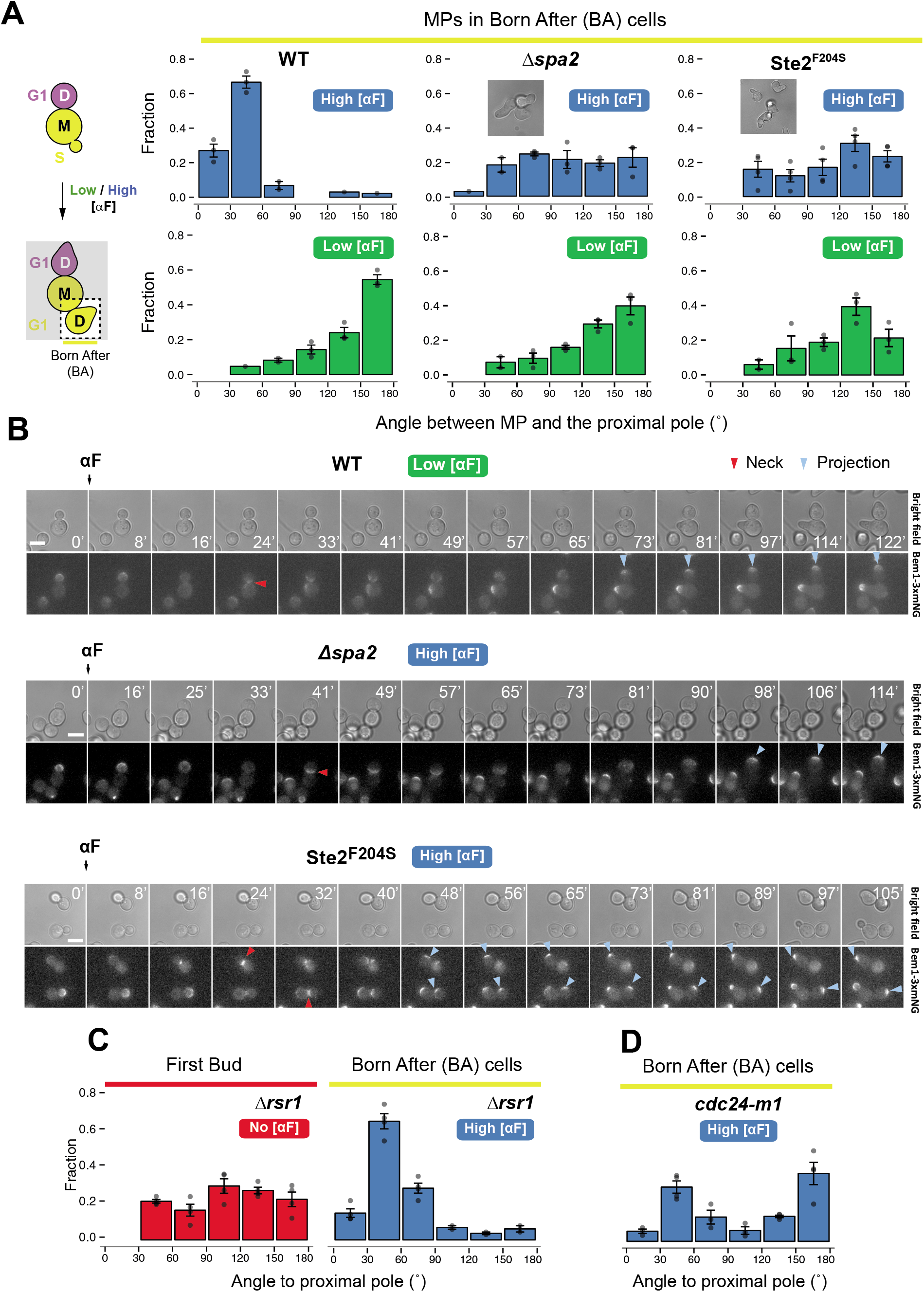
Participation of the cytokinesis-related polarity patch in MP site selection at high concentrations of α-factor. **A.** Distribution of angles of MPs relative to the proximal pole in born after cells (BA) from wild-type (ACL379), Δ*spa2* (YGV5836) and a strain co-expressing Ste2 and Ste2-F204S (YGV5841) exposed to 1μM (High [μF]) or 5 nM α-factor (Low [αF]). The fraction of cells in each 30°-bin was plotted for at least 3 independent experiments. The bar and error bars represent the mean ± SEM and the points reflect the data from each assay. **B.** Dynamics of Bem1-3xmNG followed by live cell fluorescence microscopy in wild-type cells exposed to 5nM α-factor (YGV5097) or in cell cultures stimulated with 1μM α-factor from Δ*spa2* (YGV5836) or Ste2-F204S-expressing cells (YGV5841). For all images, times of acquisition is indicated in minutes and scale bar represents 5μm. Arrows indicate the time when the polarization is at the neck or when the MP is formed. **C-D.** Angle distribution from the proximal pole of buds or MPs in cells exposed to 0 or 1 μM α-factor (No/High[αF]) in Δ*rsr1* (YGV5405) or *cdc24-m1-YFP* (YGV5832). The bars and error bars correspond to the mean ± SEM of least 3 independent experiments (points).

However, pheromone signaling does not seem to be able to maintain the patch at the proximal pole when α-factor stimulation occurs during cytokinesis. Instead, MPs are formed at greater angles relative to the proximal pole in the *bud neck* group (Figure 4B). We suspected that at this point the cell cycle has already started to inhibit the polarization at the neck. As in *Δspa2* cells, this intermediate localization may be the result of the low polarity cap mobility at high concentration α-factor that could result in MPs forming next to the inhibition zone at the neck. Indeed, lowering α-factor concentration in order to increase polarity patch wandering permitted the patch of the cells in the *bud neck* group to reach the distal pole, where the MPs finally formed (Figure S1A-B).

The above results raised the intriguing idea that the proximal default polarization site in BA cells might be independent of budding cues. We evaluated this hypothesis in *Δrsr1* cells, which bud randomly due to the interrupted communication between landmark proteins and the Cdc42 module. Indeed, *Δrsr1* BA cells form MPs in the proximal pole despite their random budding pattern (Figure 5C). This behavior was also seen in strains lacking several other budding cues, both individually and combined (Figure S2, see below for more details on the mutant tested). Thus, it seems that in this condition bud and MP site selection follow different rules. Then, we tested mother cells of BA daughters. Just like daughters, mothers of the S288C or W303 background formed a proximal MP, even in the absence of Rsr1 (Figure S1C-F).

Finally, we wondered if the proximal location of MPs in BA cells was caused by the Gβγ/Ste4-Ste18 dimer, which is already positioned at the neck at the moment of MP formation (Figure 4D). In that location, it might hold the polarity patch, for example by recruiting the Far1-Cdc24 complex. To answer that question, we studied the position of MPs in strain that expresses the *cdc24*-*m1*, an allele that codes for a protein with an impaired association with Far1 but normal binding to Rsr1. In these cells, we expected MPs to be positioned following Rsr1, as if they were budding. Indeed, we observed MPs in both poles, approximately 50% in each (Figure 5D). This result supports our hypothesis that the pheromone pathway machinery by itself is able to maintain the patch next to the bud-neck, outcompeting the positional information provided by the budding cues.

### Receptor endocytosis or phosphorylation are not involved in MP site selection

Receptor trafficking has been proposed as central for the establishment and mobility of the polarity patch (Ismael et al., 2016; Suchkov et al., 2010). Hence, we tested the role of Ste2 internalization and phosphorylation in the movement of the polarity patch away from the neck. We constructed two different Ste2 mutants. In the first one, both the ubiquitin-dependent and Sla1-dependent endocytosis pathways were impaired by replacing the 7 lysines in Ste2 C-terminal tail with arginines and by mutating the GPFAD signal to GPAAD (Ste2^7KR&GPAAD^) (Hicke and Riezman, 1996; Howard et al., 2002). In the other mutant, we also substituted twenty serines/threonines potentially phosphorylated by Yck1/Yck2 for alanines (Ste2^20STA&7KR&GPAAD^) (Ballon et al., 2006). In both cases, Ste2 variants were fused to CFP to monitor their localization (Figure 6A). Ste2^7KR&GPAAD^ could form pointed MPs and had only residual endocytosis; Ste2^20STA&7KR&GPAAD^ had no remaining detectable internalization and MPs were broader, supporting the proposed role of receptor phosphorylation in modulating polarization. Then, we analyzed the angles of the MPs in BA cells expressing these receptor variants. At high pheromone, both receptors formed proximal MPs, though with somewhat higher angles (Figure 6B). This is probably due to the failure of these receptors to polarize properly (Suchkov et al., 2010). At low pheromone, Ste2^7KR&GPAAD^ behaved as WT. In contrast, Ste2^20STA&7KR&GPAAD^ cells seemed unable to polarize away from the neck at 5 nM pheromone. However, we noticed that this mutant receptor increases the sensitivity of the strain to α-factor (Figure 6C). Therefore, 5 nM might not be a low enough concentration for this mutant strain to move the patch to the distal pole. Thus, we tested lower pheromone concentrations. Indeed, at 2 and 1 nM pheromone, Ste2^20STA&7KR&GPAAD^ mutant cells could migrate the patch from the neck and were able to reach the distal pole to form an MP (Figure 6D). Taken together these results indicate that neither receptor endocytosis nor phosphorylation are required for the selection of the polarization site during the pheromone response of cells born after pheromone stimulation.

**Figure 6.**
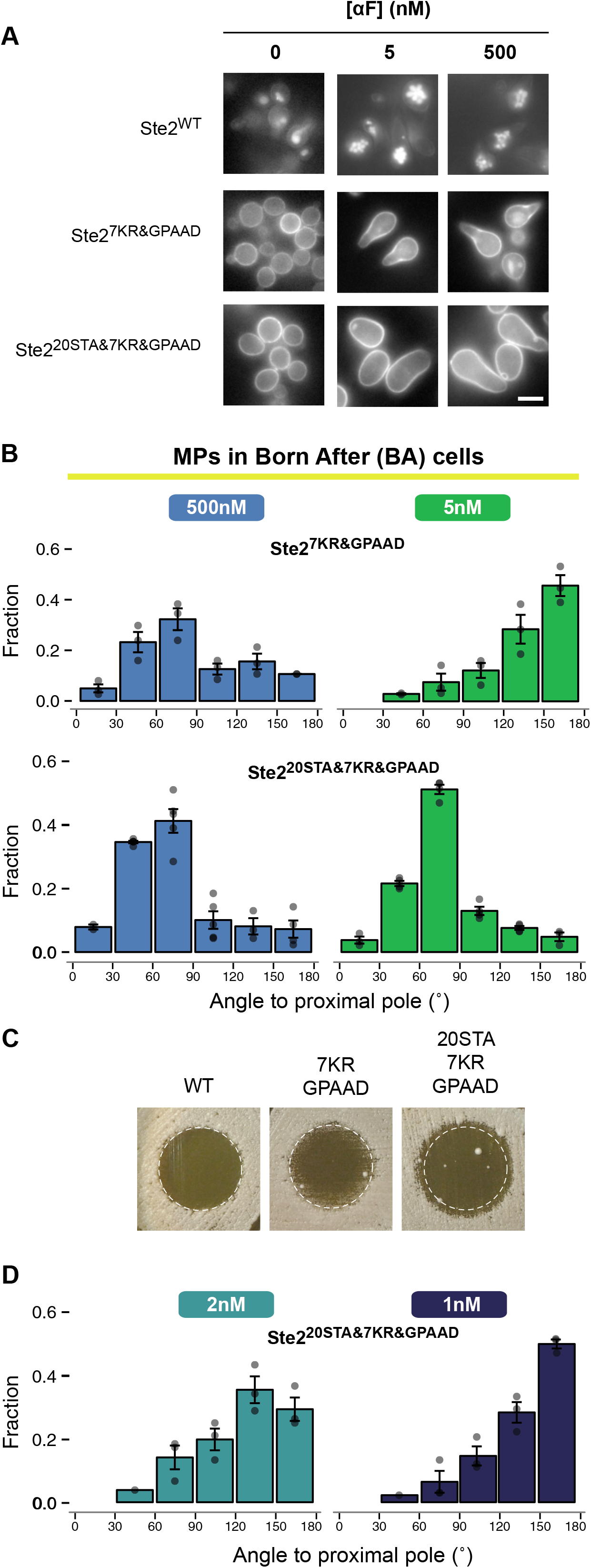
Role of receptor endocytosis and phosphorylation in mating projection default position in BA cells. **A.** Fluorescence microscopy images of *STE2* alleles C-terminally tagged with CFP exposed to 0, 5 or 1μM α-factor: Ste2^WT^-CFP (YGV5560), Ste2^7KR&GPAAD^-CFP (YGV5842), Ste2^20STA&7KR&GPAAD^-CFP (YGV5843). The scale bar represents 5μm. **B**. Distribution of MP angles from born BA cells of strains in A stimulated with 1μM (High [αF]) or 5 nM α-factor (Low [αF]). The fraction of cells in each angle bin from at least 3 independent experiments (points) is represented. The bars with error whiskers reflect the mean ± SEM. **C**. Sensitivity to pheromone of strains carrying the three *STE2* alleles was measured by Halo assays. The dashed-line circle denotes the growth arrest diameter of Ste2^WT^ cells. **D**. As in B, cells expressing Ste2^20STA&7KR&GPAAD^-CFP were exposed to 1 and 2nM α-factor and the angles between MP and the proximal pole was determined. At these doses, Ste2^WT^-CFP and Ste2^7KR&GPAAD^-CFP cells fail to form MPs.

### BB cells polarize de novo using budding cues as well as a pheromone-specific distal cue

So far, we uncovered the mechanism that promotes polarization in the proximal pole using the pre-polarized structure at the neck in a budding cue-independent manner. This mechanism applies to cells committed to the cell cycle at the time of stimulation (BA cells) with high concentrations of pheromone. But what guides the MPs into the distal pole in BA cells at low α-factor or in BB cells in high and low α-factor concentrations?

Figure 7 collects the behavior of W303 BB daughters in several deletion backgrounds exposed to high pheromone concentration. First, we analyzed the effect of inactivating the budding cues. The absence of Bud4 in W303 caused a mixture of proximal and distal budding but MPs were mostly distal (Figure 7A) indicating that the impaired axial landmark could not recruit the polarity proteins during the pheromone response. To test if Axl2 has no residual role in MP positioning, we deleted it. The resulting *Δaxl2* strain exhibited complete distal budding as well as a subtly increased use of the distal pole during MP formation (Figure 7B). This last observation suggested that the presence of proximal Axl2 affects distal polarization through a competition-based mechanism for a limiting amount of polarity proteins.

**Figure 7.**
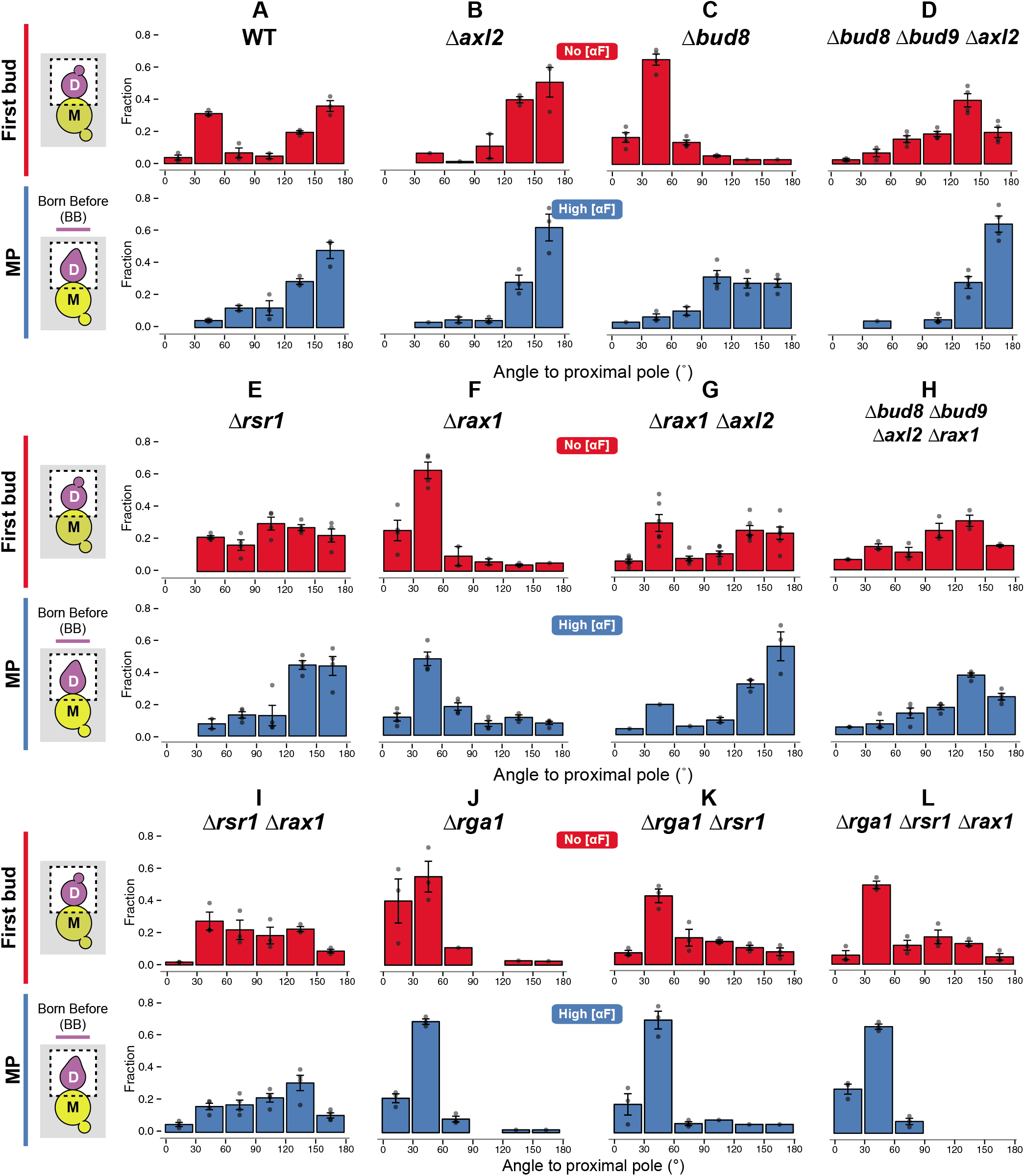
Genetic basis of MP site selection in BB cells. Distribution of angles to the proximal pole of the first bud in daughter cells (No [αF]) or of the mating projection in daughter cells born before 1μM pheromone stimulation (BB cells, High [αF]) from different mutant strains. In all cases, the angle distribution was divided in 30 degrees bins and the fraction of cells in each category was calculated. Data from at least 3 independent experiments (points) was pooled to calculate the mean ± SEM (bars and whiskers). Wild-type (WT, ACL379); Δ*axl2* (YGV5428); Δ*bud8* (YGV5429); Δ*rsr1* (YGV5405); Δ*bud8*Δ*bud9*Δ*axl2* (YGV5433); Δ*rax1* (YGV5259); Δ*rax1*Δ*axl2* (YGV5408); Δ*bud8*Δ*bud9*Δ*axl2*Δ*rax1* (YGV5839); Δ*rsr1*Δ*rax1* (YGV5838); Δ*rga1* (YGV6017); Δ*rga1*Δ*rsr1* (YGV6018) and Δ*rga1*Δ*rsr1*Δ*rax1* (YGV6035).

The natural guess for the distal landmark for MPs is Bud8, the only cue that guides bud formation at the distal pole in diploids and bipolar budding haploid variants. As expected, budding in Δ*bud8* cells was completely proximal. Strikingly, Δ*bud8* BB cells still used the distal pole for MPs, though less precisely (a somewhat broader angle distribution) (Figure 7C). Deletion of the proximal cues in this strain (Δ*bud8*Δ*bud9*Δ*axl2*) resulted in a narrower distribution of MPs around the distal pole (Figure 7D). This suggested that Bud9 and Axl2 compete for the polarization machinery, slightly destabilizing the distal polarity patch during pheromone stimulation. In light of the result of the triple cue deletion strain (Δ*bud8*Δ*bud9*Δ*axl2*), we were not entirely surprised when we found that random budding mutant Δ*rsr1* also chose the distal pole for MP formation, both in W303 and S288C (Figure 7E and S3B). Altogether, these observations pointed to the existence of a pheromone specific cue in the distal pole independent of Bud8 and Rsr1.

Next, in order to find the pheromone-specific distal cue, we evaluated the effect of deleting *RAX1* and *RAX2*.The proteins Rax1 and Rax2 are localized at both poles of the cell and together help position the bipolar landmarks Bud8 and Bud9 (Kang et al., 2004). As previously reported, the absence of Rax1 or Rax2 caused budding to become proximal, because the only remaining well positioned cue, Axl2, is proximal. Deletion of *RAX1* or *RAX2* also caused MPs to become proximal (Figure 7F and Figure S3D). This is striking since deletion of the known Rax1-Rax2 distal effector, Bud8, did not result in proximal MPs. This result suggested that the Rax1-Rax2 complex may be part of a new pheromone-promoted distal cue.

The proximal MPs in the *Δrax1* strain reverted to distal after deleting *AXL2* (Figure 7G), indicating that this cue dominated MP localization in the absence of Rax1. The distal choice in the double *Δrax1Δaxl2* was guided by Bud8, since the quadruple mutant *Δrax1Δaxl2Δbud8Δbud9* was nearly random both in terms of budding and MP formation (Figure 7H). We speculated that the slight residual distal tendency of this mutant could be dependent on Rsr1 itself. Therefore, we generated a strain deficient in both *RSR1* and *RAX1*. The double deletion *Δrsr1Δrax1* cells budded and formed MPs randomly (Figure 7I).

In summary, the above results show that the Rax1-2 complex acts, independently of the known budding cues, as a pheromone-promoted cue for distal positioning of MPs, and that Rsr1, via the classic budding cues and by a minor, cue-independent mechanism, also biases MP positioning.

Similar results were obtained at low α-factor doses both in BA and BB cells (Figure S3E and Figure S4).

### Cdc42 inhibition within birth scars is essential for correct MP localization

Recently, it was demonstrated that the Rax1-Rax2 complex interacts with the Cdc42 GAP Rga1 in the current (neck) and old cell division sites (Meitinger et al., 2014; Miller et al., 2017). In daughter cells, Rga1 remains in the proximal pole throughoutG1. This avoids re-polarization in places of weakened cell wall, such as the birth scar. In view of this, we tested if the distal localization of MPs directed by Rax1-Rax2 was in fact the reflection of Cdc42 inhibition at the proximal pole by Rga1. The fact that *Δrga1* cells showed a phenotype similar to *Δrax1*, seemed to support this idea (Figure 7J). However, *Δrsr1Δrax1* cells behaved clearly different than *Δrsr1Δrga1* cells (Figure 7K), since the latter made proximal buds. This indicated that if polarization inside the birth scar was not inhibited in daughter cells, budding could not follow budding cues and it was destined to the proximal pole. The action of Rga1 also explained the low frequency of buds in the proximal pole observed in Δ*rsr1* cells. In the case of the position of MPs, deletion of *RGA1* in all strains we tested, including the random MP strain Δ*rax1*Δ*rsr1*, resulted in proximal MP formation (Figure 7L). All in all, these results show that Rga1 has a central role in inhibiting inside birth scar polarization, and that it is essential to allow relocation of the polarity patch away from the neck. We found the same behavior in BA or BB cells at low α-factor (Figure S3E and S4).

### Interaction between gradient sensing and default polarization sites

Having defined the default polarization sites in response to α-factor, we proceeded to evaluate if these biasing mechanisms affect MP site selection in a pheromone gradient. We compared wild-type W303 and the landmark-free mutant Δ*rsr1*Δ*rax1* in artificial gradients using microfluidic devices (Ventura et al., 2014). In both strains, optimal gradient sensing was observed at a low α-factor concentration range, where, according to our results, WT daughter cells, disregarding when they were born, use the distal pole as the default polarization site (Figure 2C). Taking that into consideration, we divided the cells in three groups based on the angle between cell’s proximal-distal axis and the gradient direction (Figure 8). Group I included daughter cells whose distal pole happened to be facing the gradient source hence both internal and external cues were aligned. Compared to wild-type cells, Δ*rsr1*Δ*rax1* cells in this group oriented in the direction of the gradient less precisely (87.2±4.1% vs 59.8±2.1% of cells aligned), suggesting that, in this situation, landmarks can collaborate with gradient sensing. In Groups II and III (proximal-distal axis 90° or 180° relative to the gradient, respectively), default sites and the gradient can compete. A large faction of wild-type cells in Group II tracked the source of α-factor correctly (77.0±3.1%). Strikingly, all the remaining cells chose the distal pole (23.6±2.8%), likely following the activity of internal cues. Δ*rsr1*Δ*rax1* cells also aligned well (86.1±2.2%), and lacked the distal pole choosing group (2.2±1.5%). The most striking result was obtained in Group III, where wild-type cells tracked very poorly the direction of the gradient (18.8±4.9%), whereas Δ*rsr1*Δ*rax1* cells aligned quite efficiently (52.7±5.5%,). Note that the region directly facing the gradient (−30° to 30°) is not used. We speculate that at least two factors contributed to this. First, in this region polarization was inhibited by Rga1. Second, the presence of the mother cell might have affected the shape of the gradient, locally reducing the α-factor concentration, so that the actual perceived gradient direction was slightly shifted. Surprisingly, wild-type cells of this group did not choose the distal pole (their default site) and exhibited a rather random distribution of MP localizations instead. Thus, for this group the conflict between external and internal cues in WT cells is such that the cell’s choice is incorrect more often than not. These are the first solid evidences that indicate that intrinsic polarization landmarks can interfere with gradient sensing.

**Figure 8.**
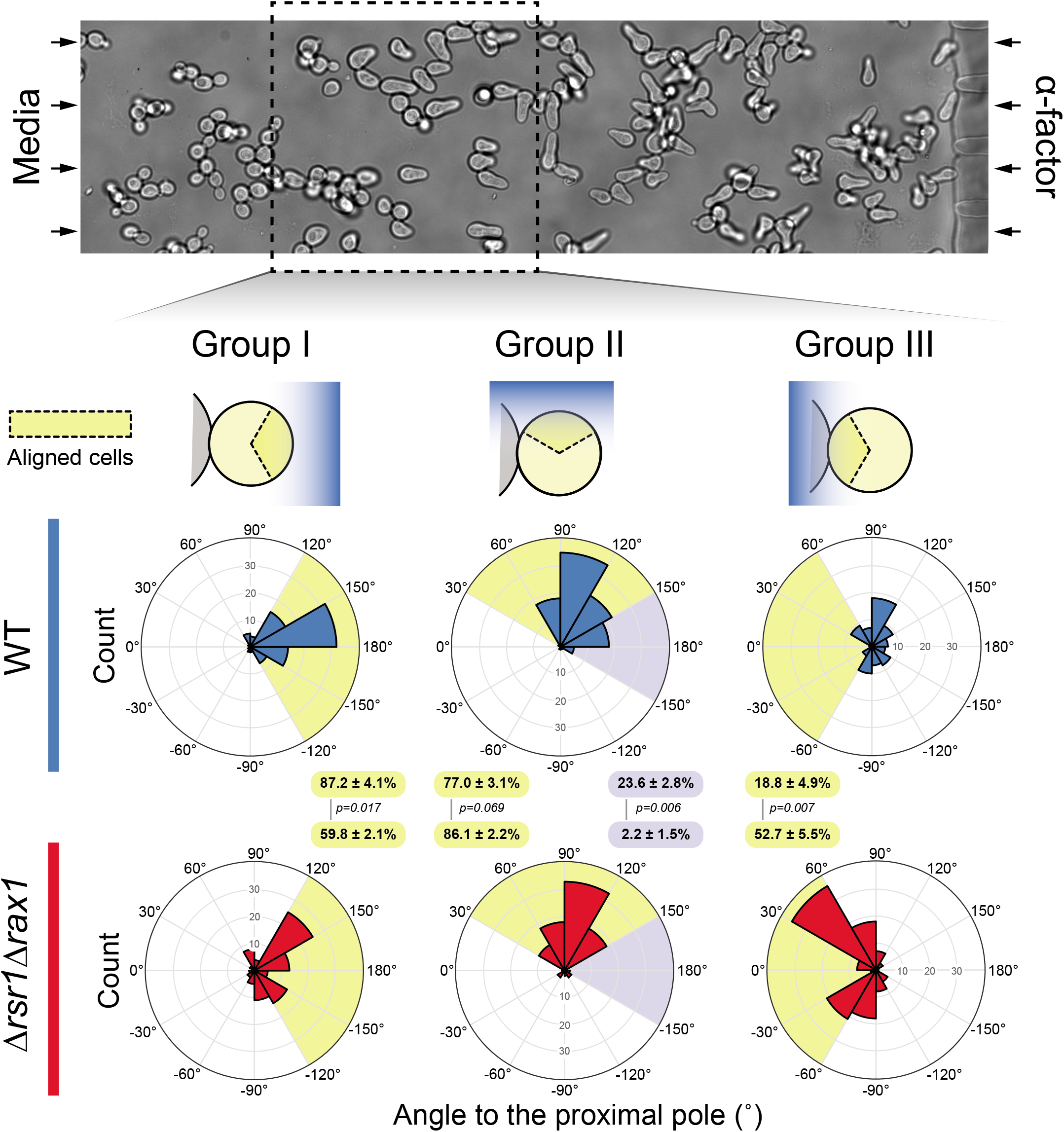
Interaction between internal polarity cues and gradient sensing. Wild-type (WT) and Δ*rsr1*Δ*rax1* strains were exposed to artificial linear gradients of α-factor generated in microfluidic devices. Above, representative image after 3hs. The box indicates the region of good cell alignment (see Materials and methods for details). Below, daughter cells were divided in 3 groups depending on the position of their distal pole relative to the gradient. In Group I, it faced the gradient (centered at 0° ranging from −60° to 60°). In Group II, the distal pole was perpendicular to the gradient (ranging from ±60° to ±120°). In Group III, the distal pole was located in opposite direction from the gradient (centered at 180° and extending from −120° to 120°). In each group, the distribution of the angle between the proximal pole and the mating projection is represented as a polar histogram (30°-bin). The data from 4 (WT) or 3 (Δ*rsr1*Δ*rax1*) independent experiments were pooled (total number of cells: 270 (WT) and 311 (Δrsr1Δrax1)). The highlighted regions correspond to the angles within each group that include cells aligned with the gradient (yellow) or the distal pole (grey). The percentage of cells in these regions is expressed as mean ± SEM. Statistical comparisons were evaluated by generalized linear models with the p-values stated.

## Discussion

In this work we revisited a matter studied since the early ’90s: the interaction between different polarization cues in yeast. Pheromone gradient sensing during mating must override the internal polarity cues used for budding. By live cell microscopy and microfluidics technics we uncovered three previously overlooked features of this signaling system. First, the cytokinesis-related polarization can serve as a polarity cue independently of all known landmark proteins. Second, the Rax1-Rax2 complex functions as pheromone specific polarity cue in the distal pole of the cells. Finally, we showed that internal cues remain active during pheromone gradient tracking and that they interfere with this process biasing the location of MPs.

### Gradient sensing and intrinsic polarity cues

Here, for the first time, we evaluated the gradient sensing ability of yeast truly devoid of intrinsic polarity cues. Contrary to budding, where deleting *RSR1* causes randomly positioned buds, MPs become randomly distributed only after deleting both *RSR1 and RAX1* (see below). Surprisingly, *Δrsr1Δrax1* cells tracked the gradient direction better than wild-type. This was especially evident when gradient and polarity cue were in opposite locations (Group III). This indicates that the constant activity of intrinsic cues does interfere with the stabilization or the localization of the polarity patch that tracks pheromone concentration. At the molecular level, this may be caused by a competition-based mechanism. Intrinsic cues may capture polarity proteins from a shared cytoplasmic pool, destabilizing a polar cap formed facing the gradient. This is similar to what happens in the resolution of multiple polarizations before budding, where one focus, via positive feedback, recruits all available polarity proteins (Wu et al., 2015). That is, the presence of a focus (in the distal pole) may partly deplete the incipient pheromone-induced patch facing the gradient, thereby destabilizing it. This may increase patch mobility, preventing a steady alignment with the gradient. Finally, through positive feedback, one particular position, more often than not away from the pheromone source, may become fixed. This idea is reminiscent of the recent proposal that patches may exist in an immature, indecisive (mobile) state, and then evolve into a committed, less mobile patch (Lew et al., 2018).

Conceptually, there is a patent antagonism between chemotropic growth and default polarization sites: gradient sensing should overcome internal cues. However, we now show that, instead of inhibiting or dominating over all pre-existing landmarks, pheromone signaling induces the utilization of specific polarity cues via Rax1-Rax2. It seems then that the use of default polarization sites could have been positively selected through evolution. In a natural environment, mating often involves germinating spores. For outcrossing to occur, yeast should polarize away from their mothers (Murphy and Zeyl, 2010). A distal bias may favor the encounter of cells from different *asci*. That is, the pheromone signaling seems to have evolved a mechanism that favors outbreeding, but, somewhat paradoxically, it interferes with efficient gradient sensing.

### MP positioning in the absence of Rsr1

Early evidence showed that wild-type yeast, when treated with pheromone, make MPs next to the proximal pole (Madden and Snyder, 1992). Disrupting bud site selection genes such as *BUD1/RSR1* and *BUD2* caused MPs to become random, leading to the idea that the default MP site is governed by the same machinery that directs bud site selection. In the light of our results, that conclusion could be a misinterpretation stemmed from the scoring method for MP localization utilized at the time. In this method, the angle between the shmoo and the position of the bud scar (in single bud scar mother cells) was measured. Because in the case of *Δrsr1* bud scars are randomly located relative to the original proximal-distal axis, MPs aligned with that axis would appear randomly located as well (see Figure S5 for a detailed explanation). Our data is consistent with previous results, but our single cell, time-lapse analysis allowed us to draw a largely different interpretation. These findings suggest that previous data obtained using Δ*rsr1* yeast as a means of severing the connection between internal cues and pheromone-induced polarization might deserve reanalysis (Deflorio et al., 2013; Dyer et al., 2013; Hegemann et al., 2015; McClure et al., 2015; Nern and Arkowitz, 1999; Strickfaden and Pryciak, 2008).

Ours is not the first evidence of distal polarization in daughter cells in the *Δrsr1* background. *Δrsr1* daughters from diploid cells in general or from haploid cells committed to invasive growth still showed a degree of distal budding (Chen and Konopka, 1996; Cullen and Sprague, 2002; Michelitch and Chant, 1996).

### Cytokinesis patch as a polarity cue

It is intriguing that the observation that the MPs are formed proximally in response to saturating α-factor even in the absence of protein landmarks has remained unnoticed for so long. The use of cytokinesis as a polarity cue may be a feature shared by many polarized cells. For example, in *D. melanogaster* during nervous system development, the mitotic cleavage site sets the axis of the precursor cell and defines where the first neurite is grown (Pollarolo et al., 2011). In epithelial cells, the mitotic contractile structure called “midbody” helps to preserve the apical–basal tissue architecture during proliferation by providing a spatial cue for the formation of the apical daughter cell interface (Herszterg et al., 2014). In yeast, polarization proximal bias enables growth of a daughter cell towards its mother after it has switched mating-type so that it mates with it; at the same time, the distal bias dependent on Rax1-Rax2 may represent a counterbalance to enable genetic outcrossing.

### A novel pheromone-promoted role for Rax1 during MP positioning

When the cytokinesis polarization is not used for MPs, yeasts employ the known budding cues or the novel pheromone-specific Rax1-Rax2 complex. Table 1 summarizes the phenotype of all mutant tested. If the Bud4-Axl2 complex is intact, like in S288C strains, MPs follow this signal in the proximal pole. In the absence of this strong cue, cells use for budding the bipolar landmarks Bud8 or Bud9, as in W303. We found that in response to pheromone, the distal Rax1-Rax2 activity may be independent of Bud8 or Bud9, as can be observed in *Δrsr1*, *Δbud8* or *Δbud8Δbud9Δaxl2*. Rax1-Rax2 dominates in W303, where the use of Axl2 is weakened by the absence Bud4. Only when the Rax1-Rax2 complex is inactivated in this background, the remaining Axl2 activity can drive polarization in the proximal pole, as may be deduced by analyzing *Δrax1* and *Δrax1Δaxl2* cells. This is similar to what happens during budding and what gave the name to Rax1: Revert to AXial (Fujita et al., 2004). Only when both budding cues and the Rax1-Rax2 complex are inactive (*Δrax1Δrsr1*) are random MPs observed. The residual distal tendency observed in *Δbud8Δbud9Δaxl2Δrax1* compared to *Δrax1Δrsr1* suggests that Rsr1 has some distal positioning activity independent of protein landmarks. In summary, in decreasing order of strength, cues for MP positioning may be sorted as follows:

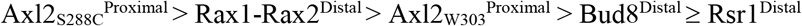

**Table 1.** Summary of budding and MP position phenotypes of mutant strains

| Strain | MP default site |  | First bud position | Genetic background |
| --- | --- | --- | --- | --- |
| | Low [αF]<br>BA&BB cells or<br>High [αF]<br>BB cells | High [αF]<br>BA cells | | |
| WT W303 | Distal | Proximal | Distal & Proximal | W303 |
| WT S288C | Proximal |  | Proximal | S288C |
| *BUD4*<sup>S288C</sup> | Proximal |  | Proximal | W303 |
| *bud4*<sup>W303</sup> | Distal |  | Distal & Proximal | S288C |
| *Δaxl2* | Distal |  | Distal | W303 |
| *Δbud8* | Distal |  | Distal | W303 |
| *Δrsr1* W303 | Distal |  | Random | W303 |
| *Δrsr1* S288C | Distal |  | Random | S288C |
| *Δaxl2Δbud8Δbud9* | Distal |  | Distal | W303 |
| *Δrax1* | Proximal |  | Proximal | W303 |
| *Δrax2* | Proximal |  | Proximal | W303 |
| *Δrax1Δaxl2* | Distal |  | Distal & Proximal | W303 |
| *Δrax1Δaxl2Δbud8Δbud9* | Distal |  | Random | W303 |
| *Δrsr1Δrax1* | Random |  | Random | W303 |
| *Δrga1* | Proximal |  | Proximal | W303 |
| *Δrga1Δrsr1* | Proximal |  | Proximal | W303 |
| *Δrga1Δrsr1Δrax1* | Proximal |  | Proximal | W303 |

As we previously discussed for gradient sensing, this orderly use of polarity cues revealed the competition-based nature of MPs site selection.

Positioning requires recruitment and activation of Cdc24, either via Rsr1, Far1 or a yet to be uncovered mechanism. Two results suggest that Rax1 operates via Ste4-Far1. First, *Δrsr1* yeast continue to make MPs distally in a Rax1-dependent manner. Second, the *cdc24-m1 Δrsr1* double mutant, in which Far1 cannot interact with Cdc24, fails to stabilize the polarity patch, moving continuously around the cell cortex (Dyer et al., 2013; Nern and Arkowitz, 2000), indicating that Rax1 is incapable of directing Cdc24 localization in the absence of Far1. In fact, Rax1 contains an RGS domain in the cytoplasmic side that binds and may regulate Gα/Gpa1 (Chasse et al., 2006). Through this interaction, Rax1 might help concentrate the G-protein in the distal pole. However, further experiments are needed to uncover the molecular players involved in Rax1 function during α-factor stimulation.

### Concentration-dependent MP site selection

Interestingly, α-factor concentration regulates the localization of MPs. Figure 9 summarizes our proposed model of MP site selection. Direct observation of the polarity patch in BA yeast revealed that only at low pheromone concentration does the patch have enough mobility to move away from this pre-polarized starting point. The dose-dependent change in patch mobility has been previously described under the “patch wandering” model (Dyer et al., 2013). In this model, vesicle delivery to a polarity patch by actin cables, instead of strengthening its position, destabilizes it, due to dilution of active polarity effectors. When this model is connected to the slow binding rate of α-factor to its receptor (Jenness et al., 1986; Ventura et al., 2014), the following scenarios emerge (McClure et al., 2015): At low pheromone concentrations, the new receptors deposited in the polarity patch remain unbound for a while and dilute the α-factor bound (active) Ste2 polar cap. In addition, unbound receptors actively inhibit G protein by recruiting the GAP Sst2 (Bush et al., 2016). This perturbation locally reduces the concentration of Cdc24, so that active Cdc42 is also reduced. Thus, the highest active Cdc42 location shifts to the side of the site of vesicle fusion, leading to movement (or wandering) of the polarity patch. In contrast, at a high α-factor concentration, binding to the receptor is fast (due to mass action) so that newly arriving Ste2 molecules are activated immediately, reinforcing the polarity cap. Our evidence supports patch wandering as the mechanism behind the dose-dependent pole choice observed.

**Figure 9.**
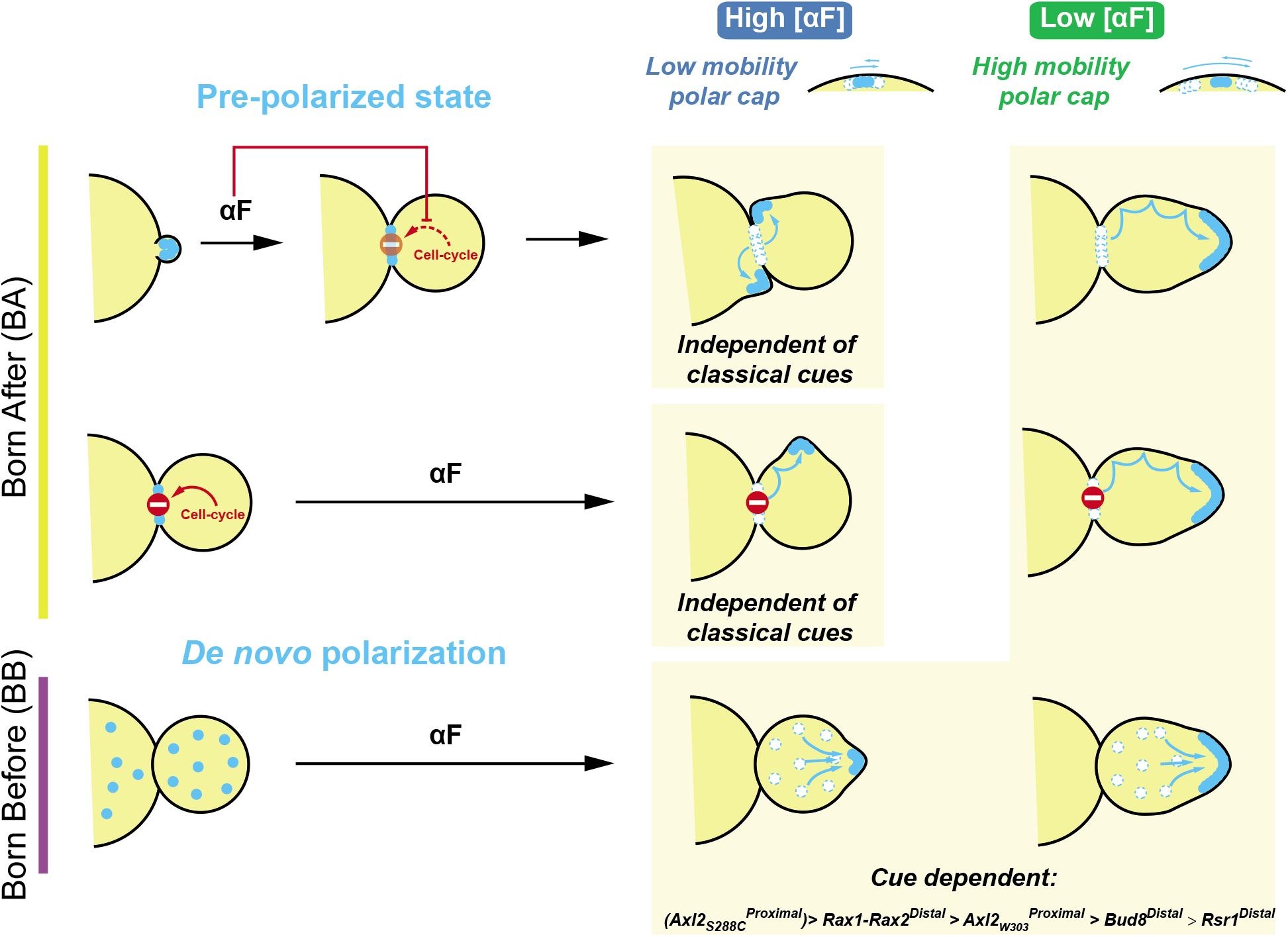
Proposed model for MP site selection. The model depicts the distinctive behavior of W303 daughter cells born after or before (BA and BB) pheromone exposure (high or low concentration). Light blue points represent the polarity patch proteins. BA cells can be further divided in two groups depending on the localization of polarity proteins at the time of α-factor stimulation. In the upper group, stimulation occurs before polarity proteins reach the neck at the endo of mitosis. In this case, at high α-factor concentrations pheromone signaling inhibits the cell-cycle dependent dispersal of the polarity patch at the neck after cytokinesis and uses it for mating projection (MP) growth. Due to the low polar cap mobility at this condition, MPs are formed in the proximal pole in a way totally independent of classical cues. For the same reason, the mothers of BA cells also form MPs in the proximal pole. At low pheromone concentrations, the polarity patch can wander around the cell cortex and then become fixed by specific cues. In W303, the distal pole is chosen through the Rsr1-dependent pathway or through a novel pheromone-specific Rax1-Rax2 mechanism. In S288C, Axl2 dominates producing MPs in the proximal pole. The second group of BA cells receives α-factor when the polarity patch is in the neck. It is likely that at this point a cell cycle-dependent mechanism is already acting destabilizing the polarization. Therefore, pheromone-induced polarization cannot occur at the proximal pole. But, given the low polar cap mobility at high α-factor, the patch cannot move long distances and MPs are finally formed laterally. The higher cap mobility observed at lower pheromone concentrations allows it to reach the distal pole as the other BA cells. In BB cells, polarization occurs de novo directly following internal cues. Based on the phenotypes of different mutants (Table 1), the order of intensity of polarity cues was established (as shown below) The diagram shows the representative behavior of W303 cells where Axl2 activity is weak.

If the pheromone pathway is activated when the polarization patch is already established at the neck, cells showed a special behavior at high pheromone concentration: MPs are placed laterally, unable to use either polar cues. To explain this different behavior, we speculate that at this point in the cell cycle, Cdc5 may have already inhibited Cdc42 at the neck (Atkins et al., 2013), making it impossible for the pheromone signaling to activate an MP at that location. When α-factor is added before this inhibition, activation of pheromone pathway may prevent this process from happening, allowing active Cdc42 to be recruited directly from the neck patch to form an MP.

Our data supports both the global and local sensing modes of gradient detection proposed in *S. cerevisiae*. Cells with a pre-polarized assembly can still correct the polar cap direction until it reaches the intrinsic cue at the distal pole in BA cells. These cells, if faced with a pheromone gradient, would correct their patch location by a local sampling mechanism. In BB cells, the polarity patch forms *de novo* in the cortical landmark. In response to a gradient, we expect them to form an MP *de novo* aligned with the gradient, by global sensing.

### Proximal to distal patch re-localization does not require receptor endocytosis or phosphorylation

Ligand-induced receptor phosphorylation and internalization has been implicated in polar cap mobility (Ismael et al., 2016; Vallier et al., 2002). However, we found no evidence supporting the role of receptor trafficking or phosphorylation in MP site selection. Strains expressing Ste2^7KR&GPAAD^, which cannot be endocytosed, or the unphosphorylatable version, Ste2^20STA&7KR&GPAAD^, still made proximal MPs at high and distal MPs at low α-factor. This results seems at odds with previous work that suggested that phosphorylation of Ste2 was necessary to switch from the proximal site to a chemotropic growth site in mating conditions (Ismael et al., 2016). However, we found that Ste2^20STA&7KR&GPAAD^ was supersensitive, thus requiring a lower α-factor concentration for the patch to reach the distal pole. Thus, in the mating mixes used in the previous work, we suspect the local concentration of α-factor was too high to allow their Ste2 allele (Ste2^7KR&GPAAD&6SA^) to wander away from the proximal site.

The increased sensitivity of Ste2^20STA&7KR&GPAAD^ is unexpected since this mutation increases the association of the receptor with the inhibitory GAP Sst2 (Ballon et al., 2006). Perhaps this mutant receptor binds ligand with more affinity or has higher GEF activity on Gα either directly or through a yet-to-be-discovered regulatory protein. In the latter case, the fraction of active G protein would be increased relative to the fraction of occupied receptors, and thus more Far1-Cdc24 complex would be recruited, potentially explaining the subsequent change in polar cap mobility. Nevertheless, a possible regulation of the pheromone pathway signaling on polar cap mobility cannot be discounted. After all, activated Fus3 is known to regulate polarity proteins, such as Bni1 and Far1 (Hegemann et al., 2015; Matheos et al., 2004).

### MP site selection in mother cells

We centered our detailed study in daughter cells to ensure the accuracy of the analysis, and tested many of the conclusions in mothers, where we found that the same principles apply. In our mid-exponential growing cultures, daughters comprise ~55% of the population and, at the time of α-factor stimulation, from the 45% of mothers, we found that 90% are committed to cell division (bud after stimulation) and only 10% are in G1 (make a MP without budding). As expected, cell cycle-committed mothers directed MPs in high α-factor concentration as BA cells: they chose the proximal pole in a cue-independent way. At lower concentrations, if mothers behave as daughters, MPs should be placed in the distal pole in a Rax1-Rax2 dependent manner. In fact, the distal polarizing activity that we observed in G1 daughter cells has also been detected in a large fraction of mother cells enriched in G1 using the *cdc28-1* temperature-sensitive allele, even in *BUD4*^+^ background (Madden and Snyder, 1992). This indicates that the mechanisms we uncovered in daughters are expected to operate in mothers as well.

### Final remarks

Most BA cells use the polarity patch from the previous round of mitosis to form MPs, which grow proximal at high α-factor concentration due to low polarity patch mobility. At low pheromone concentrations, the higher patch wandering allows the growth of MPs in the default sites also used by BB cells by *de novo* polarization. We found two signaling pathways regulating intrinsic cues: the classic Rsr1-dependent mechanism and a novel-pheromone specific branch that involves Rax1-Rax2 activity in the distal pole (Figure 9).

The presence of intrinsic polarity cues is ubiquitous in eukaryotic cells. In neurons, for example, polarity is inherited from the preexisting apical-basal polarity of the neuroepithelial progenitor cell or from the site of cell division (Yogev and Shen, 2017). These cells also must track gradients of regulatory molecules during axon guidance. The interaction between internal and external cues is, then, a common situation in cell biology. In this work, we demonstrate that intrinsic landmarks may bias the polarity system within a cell either positively or negatively depending on the situation. We expect that our results help in the understanding of gradient sensing as a cell behavior that integrate a variety of polarity systems.

## Materials and Methods

### Yeast strains

Yeast strains used in this study are listed in Table 2. Standard yeast molecular and genetic procedures were used to generate the strains. Most of the strains were generated from ACL379 which is a derivative of W303-1A (Colman-Lerner et al., 2005).

**Table 2.**
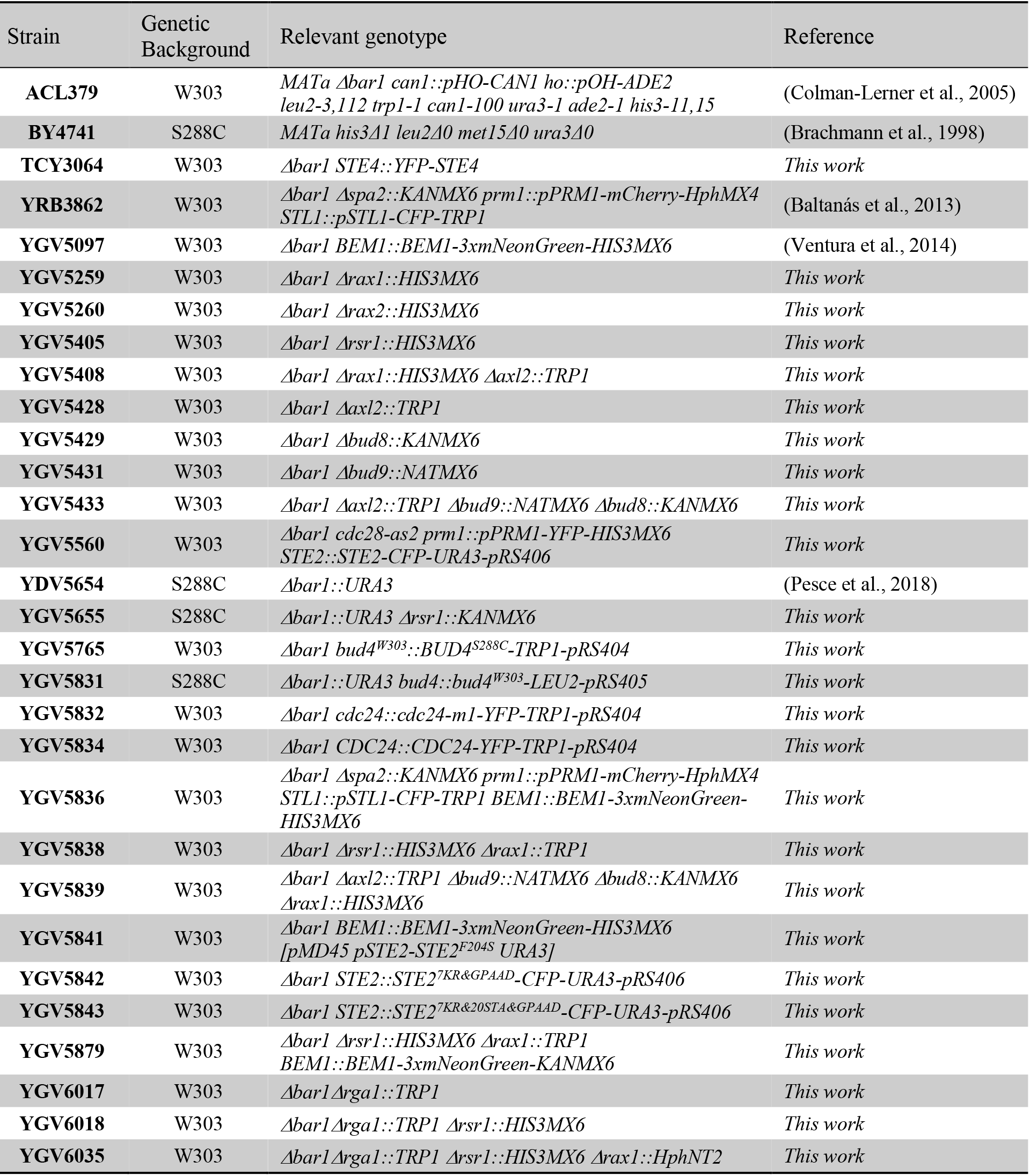
Strains used in this study.

All deletion strains were done by one-step PCR-mediated method using plasmids pFA6a-HIS3MX6, pFA6a-KANMX6, pFA6a-TRP1, pFA6a-NATMX6 or pFA6a-HphNT2 as templates (Longtine et al., 1998). In all cases, forward primers included 40nt homology arms (−40 to −1 from ATG) and the F1 sequence, and reverse oligos contained 40nt sequence following the stop codon plus the R1 primer sequence. Screening was performed by PCR and, when possible, by analyzing the budding pattern by live cell microscopy.

For tagging endogenous Bem1, also a PCR-based method was performed using the plasmid pFA6a-3xmNeonGreen-HIS3MX6 as a template (Ventura et al., 2014). Screening was made by fluorescence microscopy.

Strains carrying YFP-Ste4 were generated by integrating a plasmid containing *STE4* promoter regulating the expression of *YFP-STE4* in a pRS406 backbone. Spontaneous loop-out events were selected by plating the cells on 5-FOA (5-fluoro-orotic-acid) plates.

To express a wild-type copy of *BUD4* in our W303-derived strain, we integrated *BUD4*^*S288C*-^pRS404 plasmid linearized with StuI. This plasmid contains only the C-terminal region of BUD4 and, once integrated, results in a strain with only one functional copy of the gene. Conversely, to express the *bud4* mutant allele from W303 in S288C, *bud4*^*W303*^-*pRS405* also linearized by StuI digestion was integrated. Due to gene conversion (Rothstein, 1991), it was necessary to screen more than 40 clones by analyzing the budding pattern.

*STE2* internalization and phosphorylation defective alleles were introduced at the endogenous locus by integrating *STE2*^*WT*^*-CFP*-pRS406, *STE2*^*7KR&GPAAD*^*-CFP*-pRS406 or *STE2*^*7KR&GPAAD&20STA*^*-CFP*-pRS406 plasmids. In all cases, plasmids were linearized by ClaI digestion and screened by fluorescence microscopy.

Cdc24-m1 carrying strain was made by integrating at the endogenous *CDC24* locus the plasmid *cdc24-m1-YFP*-pRS406 digested with XcmI. Positive clones were selected based on the absence nuclear localization (Shimada et al., 2000).

To co-express the wild-type *STE2* and the *STE2*^*F204S*^ allele, which does not bind pheromone, the replicative plasmid *pMD45* was transformed in our wild-type strain (Dosil et al., 1998).

### Plasmids

Plasmids used in this work are listed in Table 3. Primers are included in Table 4.

**Table 3.**
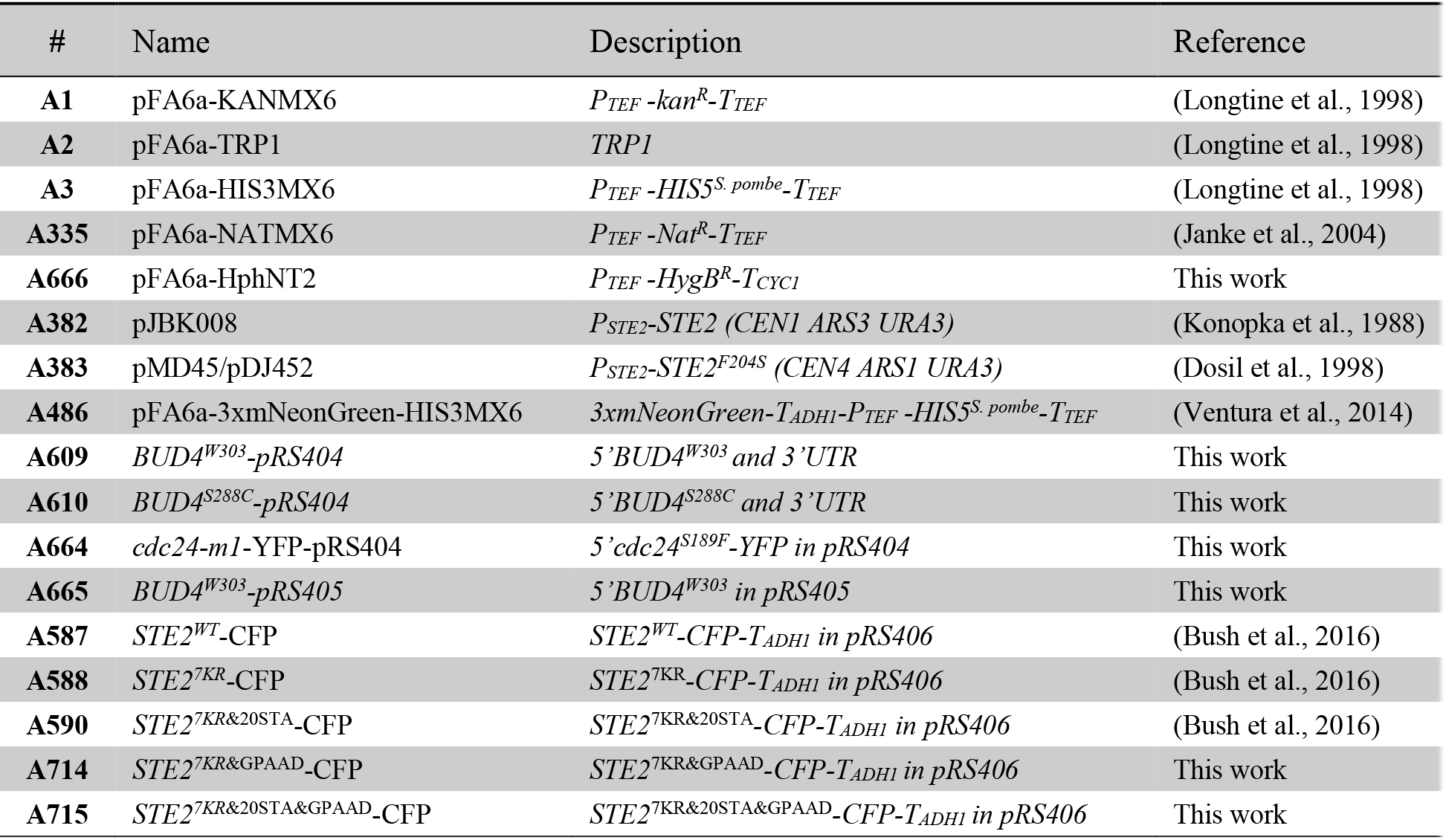
List of plasmids used or constructed.

**Table 4.**
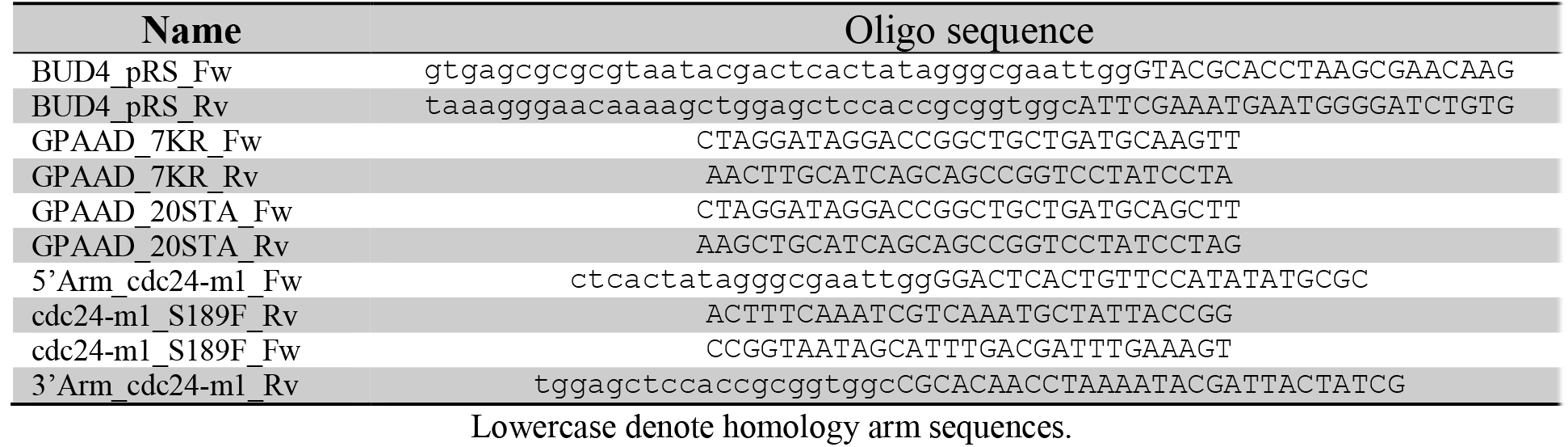
DNA sequence of oligos used for cloning

For *BUD4* cloning from W303 or S288C, a PCR fragment including the C-terminal and the 3’UTR (+2323bp for ATG to 152bp downstream the stop codon) was generated using genomic DNA from ACL379 or BY4741 as template. The primers included on both ends 35-40 nt homology arms to the pRS400 plasmid series. pRS404 was digested with NotI and Acc65I and the *BUD4* PCR fragment was cloned by isothermal assembly (Gibson et al., 2009) to generate *BUD4*^*W303*^-pRS404 and *BUD4*^*S288C*-^pRS404. To change the *TRP1* marker to *LEU2*, *BUD4*^*S288C*^-pRS404 and pRS405 were digested with BglI and the corresponding fragments were purified and ligated to form *BUD4*^*W303*^-pRS405. By sequencing the constructed plasmids, the *BUD4* mutation in W303 was confirmed: a deletion of one of the four Gs at +2456-2459 bp, counting from the ATG.

To generate plasmids carrying *cdc24*-*m1*, 2 overlapping PCR products were generated. The first one included a 20nt upstream homology arm to the pRS400 plasmid series and spanned from +119 to 581 bp of *CDC24* ORF. The second PCR product started 522 bp after ATG, continued until 201 bp after the stop codon and ended with 20nt homology arm. The S189F (+565) mutation was introduced in both internal primers. The resulting PCR products were cloned by isothermal assembly in NotI/Acc65I-digested pRS404. This plasmid was named *cdc24-m1*-pRS404. To generate the YFP tagged version of Cdc24-m1, BglII fragment from a plasmid containing *CDC24-YFP-T*_*ADH1*_ was ligated in the BglII site at +2235 of *CDC24* ORF in *cdc24-m1*-pRS404.

Ste2 mutant alleles were constructed based on our previous Ste2^7KR^-CFP and Ste2^7KR&20STA^CFP plasmids (Bush et al., 2016) by site-directed mutagenesis using the primers listed in Table 4. In both cases, the GPFAD endocytosis signal was mutated to GPAAD to block the Sla1-dependent endocytosis.

### Time-lapse microscopy methods

Cultures in mid-exponential growth were mildly sonicated to disperse clumps and diluted to a final concentration of 2.5×10^5^ cell/mL. Polyethylene glycol (PEG, avg MW=3550, Sigma) 0.1% was added to all media to avoid nonspecific binding of α-factor to plastic (Liu et al., 2013). Then, 20μL of cell suspension was applied to 384-well glass-bottom plates that had been pre-coated with 1 mg/mL concanavalin A type V (Con A, Sigma). Plates were centrifuged at 600 r.p.m. to assist the attachment of cells to the glass. Once in the microscope, 2-3 image fields per well were manually selected and the time-lapse imaging was started. After the first time point, 20μL of α-factor (Anaspec, Inc.) solutions were added to final concentrations ranging 0 to 1000nM depending on the experiment.

For imaging, an Olympus IX-81 microscope, with an Olympus UplanSapo objective with a magnification of 63× (N.A. = 1.35) coupled with an HQ2 (Roper Scientific) cooled CCD camera was used. Illumination was provided by a CoolLED pE-2 system with different filter sets for YFP and CFP (41028 and 31044v2, Chroma Technologies Corp.). The microscope was also equipped with a motorized XYZ stage and ZDC autofocus that allowed automatic field tracking. For assessing Bem1-3xmNG or YFP-Ste4 localization, Z-stacks were acquired and maximum fluorescence projections were used for visualization.

### Chemotropism assay in microfluidic devices

Microfluidic devices designed for the generation of stable gradients in open chambers were fabricated using standard protocols for polydimethylsiloxane (PDMS) microfluidic device construction as previously described (Keenan et al., 2006; Ventura et al., 2014). Briefly, silicon molds were generated by three-layer SU-8 photolithography, which were then used for making PDMS replicas of the device, by excluding PDMS from the tallest features of the mold, thereby producing open (roofless) chambers. After polymerization, the patterned PDMS structure was peeled off and bonded it onto glass cover slides by plasma-oxygen treatment (660 mtorr, 60 W, 60 s).

To improve adherence of cells to the glass, the bottom of the chambers was treated with poly-D-lysine (1 mg/mL; Sigma) at 4°C overnight and then incubated with ConA (1 mg/mL; Sigma) for 1 hour at room temperature. Next, the device was filled with 0.22μm of sterile, filtered water using a vacuum-assisted method (Monahan et al., 2001). Subsequently, the two ports of the device were connected with tubing and syringes filled with filtered synthetic complete medium alone or with 50-150nM αF and 0.1 mg/mL bromophenol blue (BPB) as tracking dye. All media contained 0.1% w/v PEG3000 (Sigma) to prevent nonspecific αF binding to the container’s surfaces (Liu et al., 2013). Water hydrostatic pressure drove all flow. Formation of the gradients was monitored by measuring BPB fluorescence. Finally, the flow was stopped, the chambers were washed with media, and a mildly sonicated yeast exponential culture was loaded on top of the device. Cells were allowed to settle and bind to the bottom glass before resuming the flow. Imaging was performed as described for glass bottom plates.

### Angle determination

Angles were manually measured using custom-written macros for ImageJ on brightfield time-lapse stacks. As explained in the main text, daughter cells were classified in born before (BB) or born after (BA) cells relative to the α-factor stimulation. A daughter cell was considered as BB only if its mother was already committed to a new round of cell division (ie, budded in the presence of pheromone). This ensured that the new born daughter was already in G1. BA cells comprised daughter cells that were buds at the time of pheromone exposure. By analyzing Bem1-3xmNG localization, BA cell could be further categorized in 3 groups: *incipient bud*, *bud periphery* and *bud neck* as in Figure 4B. However, in all other figures, BA cells are defined using bright field images. In this case, BA cells were classified as such only if we measured an increase in size between the first two time points, an unequivocal sign that that bud is still a bud, and not a new daughter. This method, however, discards cells with the patch at the bud neck, since their growth is not detectable by light microscopy. As a consequence, our classification of BA cells based on morphology alone excludes the *bud neck* group.

### Halo Assays

100 μL of mid-exponential growing cultures at 10^6^ cell/mL were plated on YPD plates. Then, 0.1 nmoles of α-factor (5μL of 20μM) were carefully placed in the middle of the plate. The plate was incubated for 2-3 days at 30°C until the zone of growth arrest became evident.

### Data analysis

All data analysis was done in R (R-Development-Core-Team, 2008). For angle histograms, cells were split into 6 bins, from 0 to 180°, according to the position of the mating projection, and the fraction of cells in each bin, in each experiment was calculated. The mean ± SEM of at least 3 independent assays was plotted. For gradient experiments, first the region of the device where cells could detect gradient direction was established. To do that, the chambers were divided in 50μm windows parallel to the gradient direction. In each of them, the standard deviation of the angle of alignment was calculated. In a region of good gradient sensing, the dispersion of angles drops below the level of the distribution of angles in cell populations exposed to isotropic α-factor. These values were used to set a dispersion-based cut-off to define the region of the devices used for analysis. In these experiments, only the first polarization was taken into account and 2 angles were scored: the angle of the mating projection (θ) and the distal pole (α) both relative to the gradient direction. Based on the second angle, the cells were divided in three groups: distal pole facing the gradient (I, −60°<α<60°); distal pole perpendicular to the gradient (II, 60°<α<120° and −60°>α>-120°); and proximal pole facing the gradient (III, 120<α and α>-120°). The angle of MPs relative to the proximal pole was determined using θ and α and, considering as positive angles the hemisphere of the cell that is facing the gradient. For Groups I and III this direction is CW. Group II is subdivided: if 60°<α<120° the direction is CW and, if −60°>α>-120°, CCW. In each group, a polar histograms with 30°-bins were plotted pooling data from 3 (WT) of 4 (*Δrsr1Δrax1*) independent microfluidic assays.

Statistical difference in the percentage of aligned cells was calculated using a generalized linear model with binomial distribution (logit link function) of the proportions of aligned cells from different experiments.

## Acknowledgments

We thank Peter Pryciak, Pablo Aguilar, Andreas Constantinou and Paula Dunayevich for helpful discussions and comments on the manuscript; Albert Folch lab for assistance and reagents for microfluidic experiments; Andreas Constantinou for providing plasmid A666.

Work was supported by Grant PICT2013-2210, PICT2015-3824, PICT2016-0949 from the Argentine Agency of Research and Technology (to A.C.L.) and Grant 1R01GM097479-01 from the National Institute of General Medical Sciences, National Institutes of Health (to Roger Brent and A.C.L.).

## Authors contributions

G.V. and A.C.L. design research. G.V. performed experiments and analyzed data. G.V. and A. C.L. wrote the paper.

---

**Figure S1.**
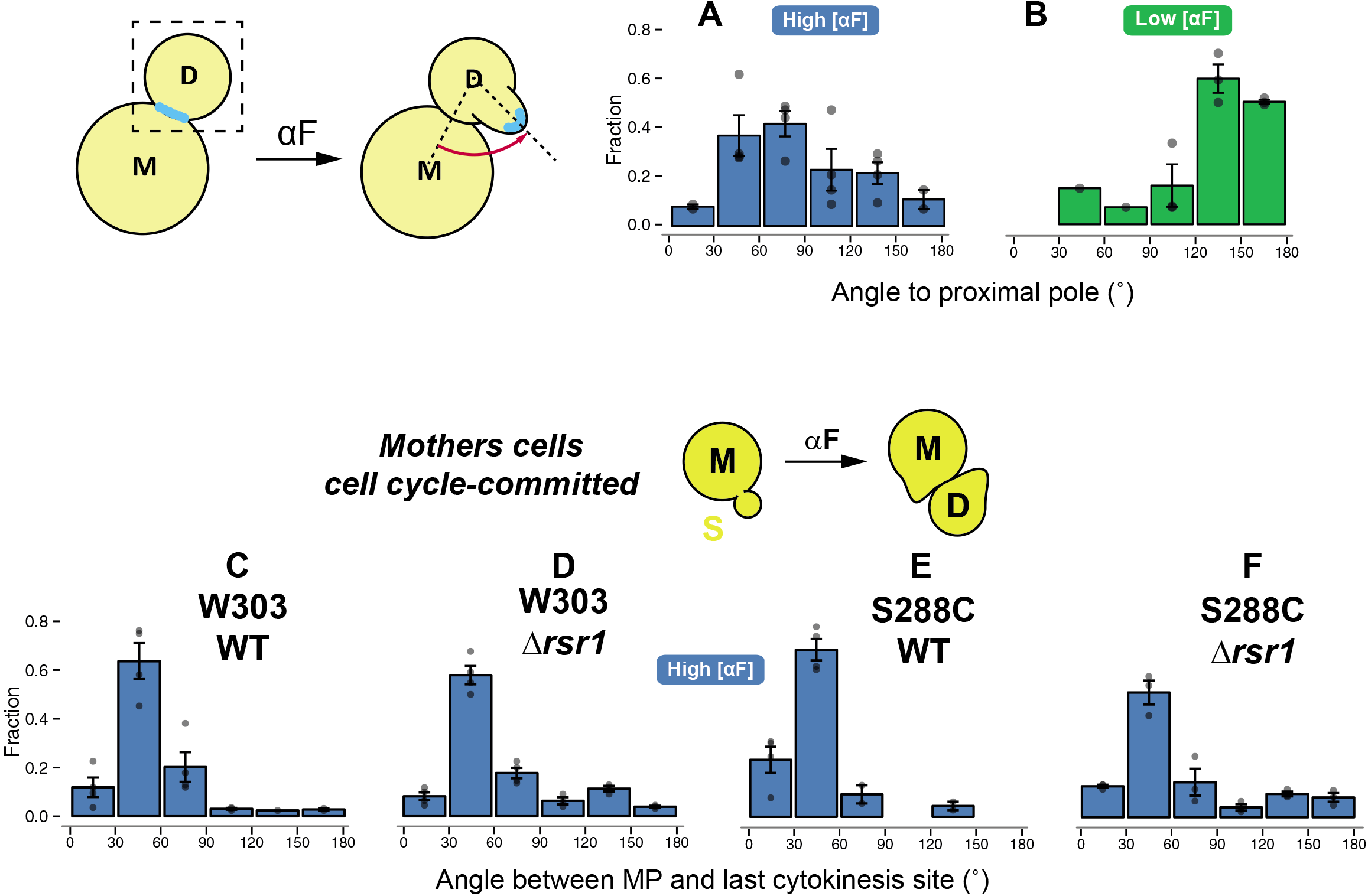
MP site selection in BA cells stimulated at the time of cytokinesis and in mother cells. **A-B.** Distribution of MPs angles in cells stimulated with low (5nM) or high (1μM) α-factor concentrations at the time of cytokinesis as judged by Bem1-3xmNG localization (Bud-neck group). The fraction of cells in each angle bin from at least 3 independent experiments (points) is represented. Strain: YGV5097. **C-F**. The angle between MP of mothers of BA cells was measured relative to the site of last cytokinesis, i.e. the place of the BA daughter cell. The angle distribution of W303 Δ*bar1* (ACL379), S288C Δ*bar1* (YDV5654), W303 Δ*bar1* Δ*rsr1* (YGV5405) and S288C Δ*bar1* Δ*rsr1* (YGV5655) cells exposed to 1μM pheromone (High [αF]) are show from at least 3 independent experiments.

**Figure S2.**
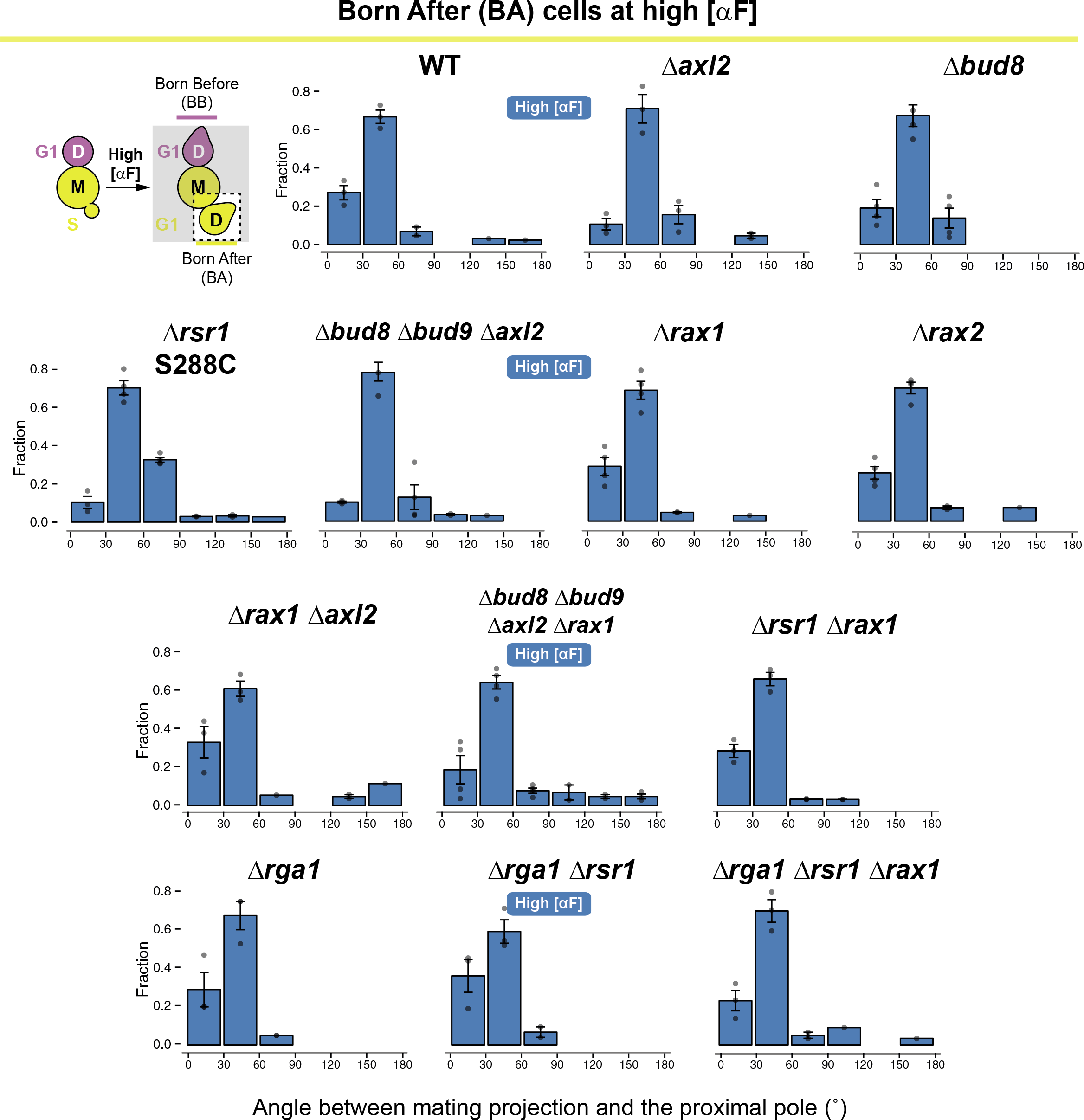
Behavior BA cells at high α-factor concentration stimulation in different mutants. Distribution of angles to the proximal pole of the mating projection in daughter cells born after 1μM pheromone stimulation (BA cells, Low [αF]) from different mutant strains. In all cases, the angle distribution was divided in 30 degrees bins and the fraction of cells in each category was calculated. Data from at least 3 independent experiments (points) was pooled to calculate the mean ± SEM (bars and whiskers). Wild-type (WT, ACL379); Δ*axl2* (YGV5428); Δ*bud8* (YGV5429); Δ*rsr1* (YGV5405); S288C Δ*bar1* Δ *rsr1* (YGV5655); Δ*bud8*Δ*bud9*Δ*axl2* (YGV5433); Δ*rax1* (YGV5259); Δ*rax2* (YGV5260); Δ*rax1*Δ*axl2* (YGV5408); Δ*bud8*Δ*bud9* Δ*axl2*Δ*rax1* (YGV5839); Δ*rsr1*Δ*rax1* (YGV5838); Δ*rga1* (YGV6017); Δ*rga1*Δ*rsr1* (YGV6018) and Δ*rga1*Δ*rsr1*Δ*rax1* (YGV6035).

**Figure S3.**
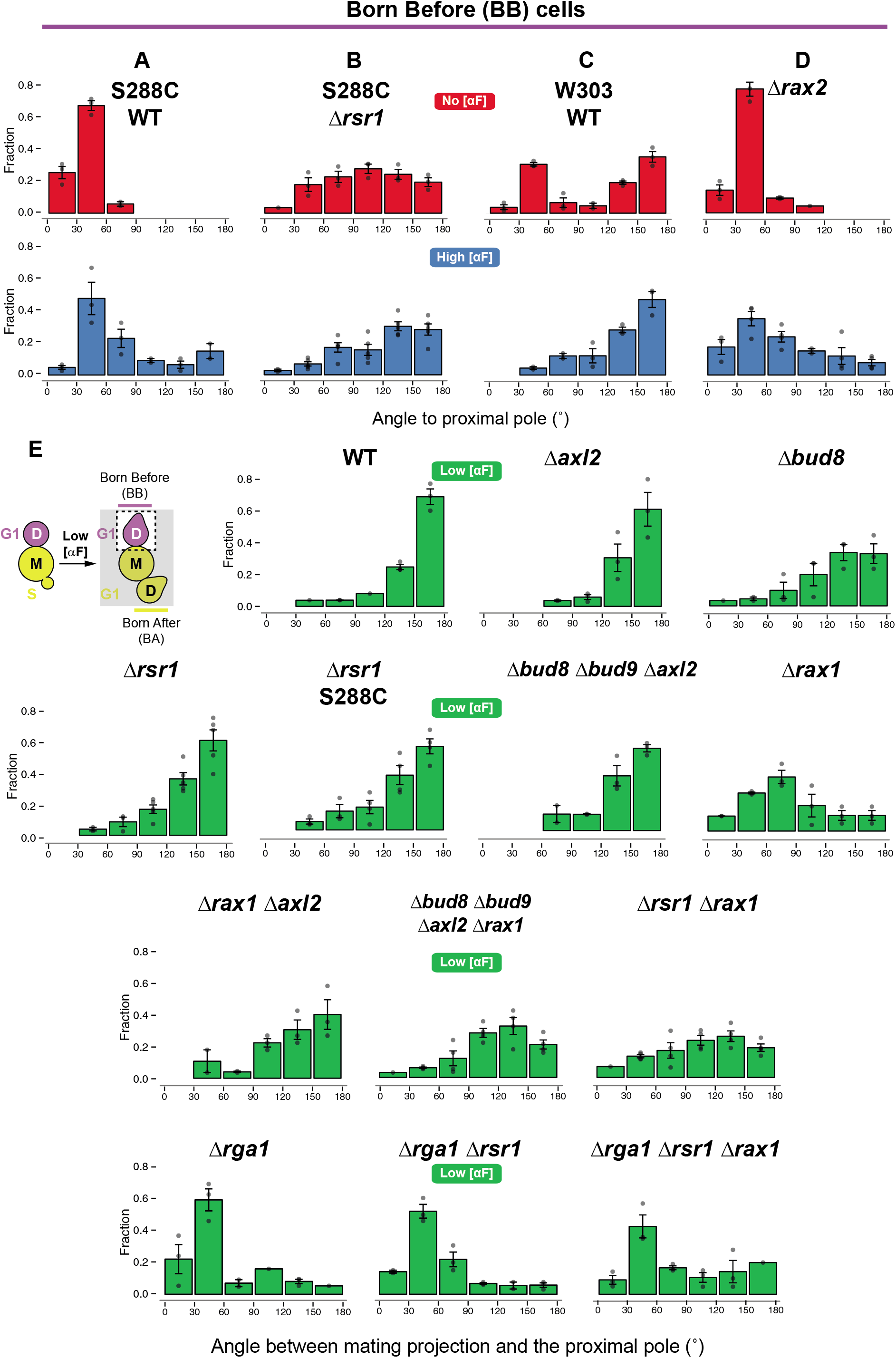
MP site selection of BB cells mutant strains. **A-D**. Distribution of angles to the proximal pole of the first bud (No [αF]) or of the mating projection in daughter cells born before 1μM pheromone stimulation (BB cells, High [αF]) from the following mutant strains: S288C Δbar1 (WT, YDV5654), S288C Δbar1 Δrsr1 (YGV5655) W303 Δbar1 (WT, ACL379) and Δrax2 (YGV5260). In all cases, the angle distribution was divided in 30 degrees bins and the fraction of cells in each category was calculated. Data from at least 3 independent experiments (points) was pooled to calculate the mean ± SEM (bars and whiskers). **E**.Distribution of angles to the proximal pole of the mating projection in daughter cells born before 5nM pheromone stimulation (BB cells, Low [αF]) from different mutant strains. In all cases, the angle distribution was divided in 30 degrees bins and the fraction of cells in each category was calculated. Data from at least 3 independent experiments (points) was pooled to calculate the mean ± SEM (bars and whiskers). Wild-type (WT, ACL379); Δ*axl2* (YGV5428); Δ*bud8* (YGV5429); Δ*rsr1* (YGV5405); S288C Δ*bar1* Δ*rsr1* (YGV5655); Δ*bud8*Δ*bud9*Δ*axl2* (YGV5433); Δ*rax1* (YGV5259); Δ*rax1*Δ*axl2* (YGV5408); Δ*bud8*Δ*bud9*Δ*axl2*Δ*rax1* (YGV5839); Δ*rsr1*Δ*rax1* (YGV5838); Δ*rga1* (YGV6017); Δ*rga1*Δ*rsr1* (YGV6018) and Δ*rga1*Δ*rsr1*Δ*rax1* (YGV6035).

**Figure S4.**
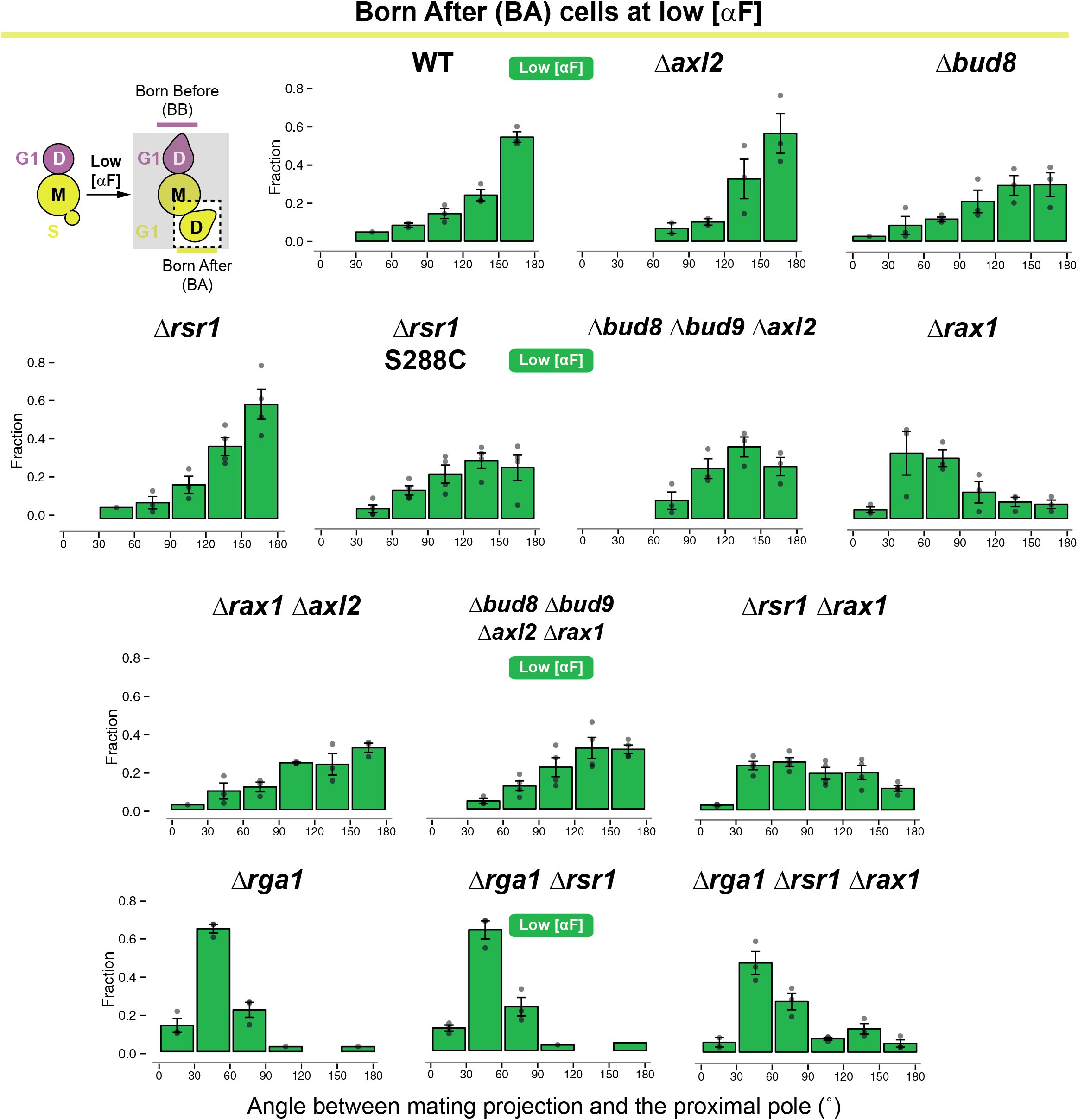
Behavior BA cells at low α-factor concentration stimulation in different mutants. Distribution of angles to the proximal pole of the mating projection in daughter cells born after 5nM pheromone stimulation (BA cells, Low [αF]) from different mutant strains. In all cases, the angle distribution was divided in 30 degrees bins and the fraction of cells in each category was calculated. Data from at least 3 independent experiments (points) was pooled to calculate the mean ± SEM (bars and whiskers). Wild-type (WT, ACL379); Δ*axl2* (YGV5428); Δ*bud8* (YGV5429); Δ*rsr1* (YGV5405); S288C Δ*bar1* Δ *rsr1* (YGV5655); Δ*bud8*Δ*bud9*Δ*axl2* (YGV5433); Δ*rax1* (YGV5259); Δ*rax1*Δ*axl2* (YGV5408); Δ*bud8*Δ*bud9*Δ*axl2*Δ*rax1* (YGV5839); Δ*rsr1*Δ*rax1* (YGV5838); Δ*rga1* (YGV6017); Δ*rga1*Δ*rsr1* (YGV6018) and Δ*rga1*Δ*rsr1*Δ*rax1* (YGV6035).

**Figure S5.**
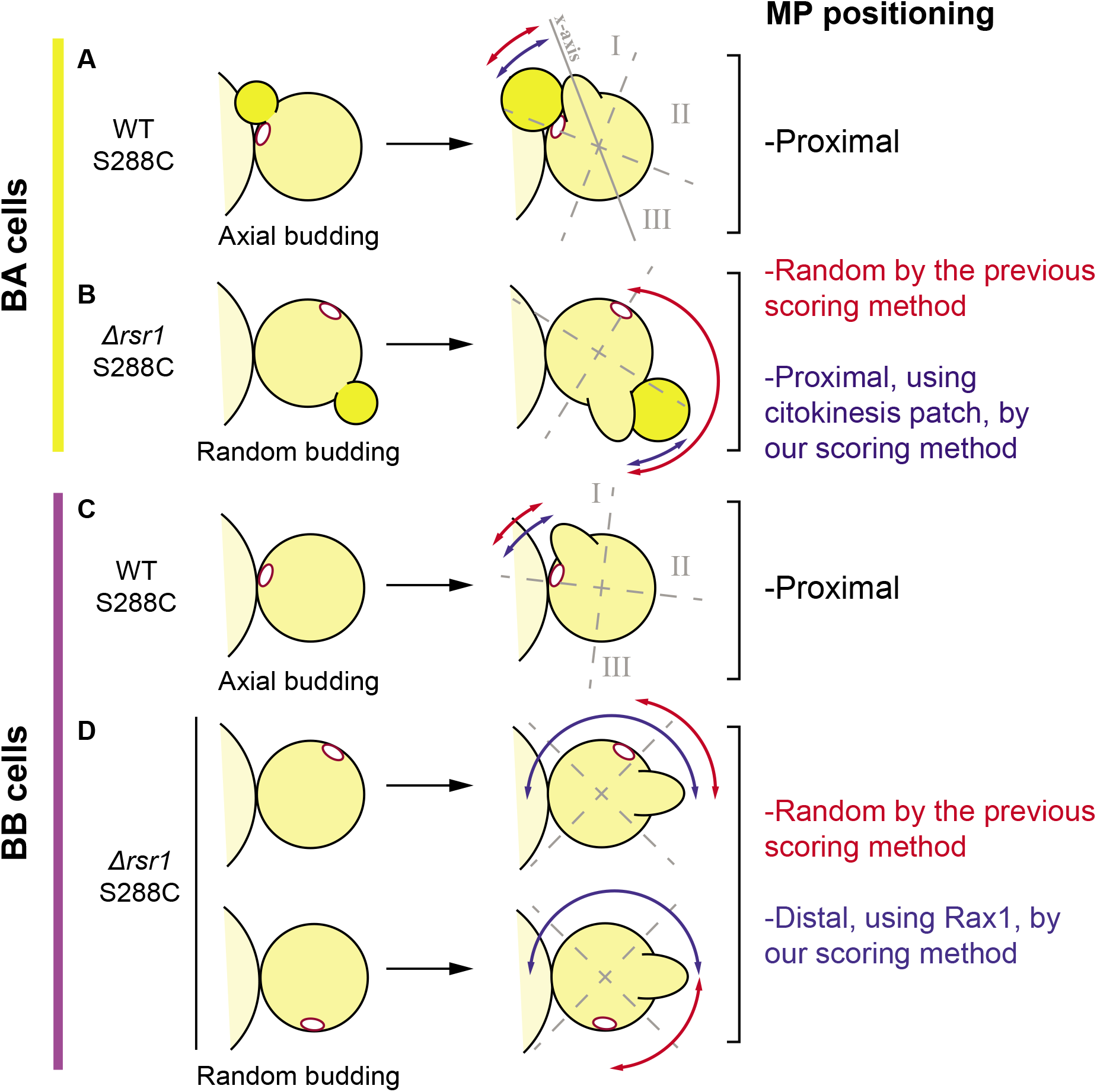
Re-interpretation of original data from Madden and Snyder (1992). In the original work, MP positioning was scored on cells with only one bud scar as revealed by calcofluor white staining. In this diagram we present these cells next to their mother to highlight the position of the proximal pole/birth scar. The authors quantified the distribution of sites selected for mating projection formation by dividing each cell into three distinct domains. These domains were determined relative to an x-axis running the length of the cell specified by the projection. Class I cells contained bud scars located −45 to +45° from the x-axis at the end of the cell containing the projection. Class III cells contained bud scars −45 to +45°from the horizontal axis on the other side of the cell. Class II cells contained bud scars in the center. The boundary between classes is indicated by dashed lines. Based on this classification we considered the results we would expect based on our current knowledge of the system. **A**. In WT BA cells, given the high α-factor concentration used, MPs are expected to form next to the cytokinesis site. In the S288C background, the budding pattern is axial and then, the previous daughter (and bud scar) would be in the proximal pole. Class I category is expected to be the most populated. **B.** However, in a Δ*rsr1* background the scenario is different. Based on our data, MPs are formed, again, next to the last daughter in a cue-independent way. Under our scoring, this would be proximal MPs. But, in the original analysis, the previous bud scar was used as reference. Given the random budding phenotype, this reference is randomly located relative to the subsequent daughter whose cytokinesis patch served as seed for MP growth. So, the random MP site selection in the cells is merely a consequence of random budding and not of random positioning of the MP. **C.** A similar outcome to A is thought to occur in WT BB cells but, in this case, by de novo polarization, MPs would form next to the last bud scar guided by Axl2 activity. In WT cells, our results and the original data concur. **D.** In Δ*rsr1* BB cells, we expect MP to form in the distal pole of the cell following Rax1. Note that the last bud scar is randomly located relative to the distal-proximal axis of the cell in this strain. Thus, as for BA cells, MPs position was measured using a random reference and therefore, MP distribution was incorrectly defined as random. The misinterpretation of the data explains why our model for MP site selection went unnoticed.

